# Arabidopsis root lipid droplets are hubs for membrane homeostasis under heat stress, and triterpenoid synthesis and storage

**DOI:** 10.1101/2025.03.24.644787

**Authors:** Patricia Scholz, Janis Dabisch, Alyssa C. Clews, Philipp W. Niemeyer, Ana C. Vilchez, Magdiel S. S. Lim, Siqi Sun, Lea Hembach, Fabienne Dreier, Katharina F. Blersch, Lea M. Preuß, Martin Bonin, Elena Lesch, Yuya Iwai, Takashi L. Shimada, Jürgen Eirich, Iris Finkemeier, Katharina Gutbrod, Peter Dörmann, You Wang, Robert T. Mullen, Till Ischebeck

**Affiliations:** University of Göttingen, Albrecht-von-Haller-Institute for Plant Sciences, Department of Plant Biochemistry, 37077 Göttingen, Germany; ENS Lyon – Laboratoire Reproduction et Développement des Plantes, 69364 Lyon, France; University of Münster, Institute of Plant Biology and Biotechnology (IBBP), Green Biotechnology, 48143 Münster, Germany; University of Guelph, Department of Molecular and Cellular Biology, Guelph, ON N1G 2W1, Canada; Chiba University, Graduate School of Horticulture, Matsudo648, Matsudo-shi, Chiba, 271-8510, Japan; Plant Molecular Science Center, Chiba University, Chiba-shi, Chiba, 260-8675, Japan; Research Center for Space Agriculture and Horticulture, Chiba University, Matsudo-shi, Matsudo, 271-8510, Japan; University of Münster, Institute of Plant Biology and Biotechnology (IBBP), Plant Physiology, 48149 Münster, Germany; Institute of Molecular Physiology and Biotechnology of Plants (IMBIO), University of Bonn, Bonn, Germany

**Keywords:** lipid droplets, roots, heat stress, proteomics, Arabidopsis, triterpenes

## Abstract

- Plant lipid droplets (LDs) and their associated proteins have numerous subcellular and physiological functions. While considerable progress has been made for LDs in many tissues, the function and composition of LDs in roots remains largely unexplored.
- We investigated the changes of the number of LDs and of the lipidome in heat-stressed *Arabidopsis thaliana* roots. Furthermore, we isolated root LDs from the Arabidopsis mutant *trigalactosyldiacylglycerol 1-1 sugar dependent 1-4* and investigated their proteome and lipidome.
- Heat stress lead to a degradation of membrane lipids and an increase in TAGs and LDs. while, fatty acid SEs decreased, probably acting as precursors for acylated sterol glycosides. A variety of proteins were enriched in root LDs, which are thus far not described as LD proteins. Transient expression of these proteins in many cases confirmed their LD localization, for example of the triterpene biosynthetic enzymes thalianol synthase and marneral synthase. We could furthermore show that the educts and products of these enzymes are enriched in root LDs, too.
- We conclude that root LDs simultaneously act as a sink and source during heat stress-induced membrane remodeling. Furthermore, root LDs play a pivotal role in triperpene synthesis and storage, thereby highlighting LDs as hubs in specialized metabolism.

## INTRODUCTION

Lipid droplets (LDs) are subcellular structures that store hydrophobic molecules such as triacylglycerols (TAGs) and steryl esters (SE) in their core and are surrounded by a phospholipid monolayer with various associated proteins (Bouchnak *et al*., 2023; Guzha *et al*., 2023). LDs are formed at the endoplasmic reticulum (ER) and protrude into the cytosol (Scholz *et al*., 2022). However, whether they actually fully detach from the ER, is not clear. In land plants, LDs are most abundant in seeds, spores and pollen (Guzha *et al*., 2023), but are also present in root and leaf tissues (Kelly *et al*., 2013; Pyc *et al*., 2017). In seeds and spores, one key role of TAG stored in LDs is to provide energy and carbon for cellular growth in the absence of photosynthesis and without provision of sugars from other organs and tissues (Huang *et al*., 2009; Zienkiewicz & Zienkiewicz, 2020; Hembach *et al*., 2024), (Turesson *et al*., 2010; Niemeyer *et al*., 2022). Similarly, in pollen tubes, LDs are thought to function as an energy and carbon source (Zienkiewicz *et al*., 2013), although they can also act as a sink for membrane lipid-derived acyl-chains during heat-induced membrane remodeling (Krawczyk *et al*., 2022a). Likewise in leaves, TAG-filled LDs act as a sink and can accumulate under different abiotic stresses (Mueller *et al*., 2015; Gidda *et al*., 2016; Doner *et al*., 2021), while membrane lipids are remodeled (Higashi *et al*., 2015; Tarazona *et al*., 2015; Shiva *et al*., 2020; Scholz *et al*., 2024). Furthermore, recent work has implied that leaf LDs play a role in pathogen defense (Hanano *et al*., 2015; Shimada *et al*., 2015; Fernández-Santos *et al*., 2020) and are important for stomatal development (Ge *et al*., 2022). Recently, LDs have been also associated with the synthesis of furan-containing fatty acids (Omata *et al*., 2024).

Given the array of functions of LDs across different plant tissues, it is not surprising that their proteomes are also highly diverse (Horn *et al*., 2013; Brocard *et al*., 2017; Kretzschmar *et al*., 2018; Fernández-Santos *et al*., 2020; Kretzschmar *et al*., 2020; Doner *et al*., 2021; Niemeyer *et al*., 2022; Hembach *et al*., 2024; Omata *et al*., 2024; Scholz *et al*., 2024) and can display marked changes during stress (Scholz *et al*., 2024) and developmental processes (e.g., seedling establishment, Kretzschmar *et al*., 2020; and spore germination in the moss *Physcomitrium patens*, Hembach *et al*., 2024). For example, oleosins act as major LD surface proteins in desiccation tolerant tissues such as seeds (Huang, 2018), pollen (Roberts *et al*., 1993), spores (Huang *et al*., 2009), and the tubers of yellow nutsedge (*Cyperus esculentus*) (Niemeyer *et al*., 2022), but are notably absent in leaves (Brocard *et al*., 2017; Scholz *et al*., 2024). However, oleosins have recently also been found as the major LD proteins in the streptophyte alga *Mesotaenium endlicherianum* when nutrient stressed but fully hydrated (Dadras *et al*., 2023). Even more specific to seeds seem to be the proteins LIPID DROPLET PROTIEN OF SEEDS (LDPS) and SEED LIPID DROPLET PROTEIN (SLDP) (Kretzschmar *et al*., 2020), with the latter and its interaction partner LIPID DROPLET PLASMA MEMBRANE ADAPTOR (LIPA) mediating a membrane contact site between LDs and the plasma membrane (Krawczyk *et al*., 2022b). Other notable plant LD proteins are more ubiquitously found. These include LIPID DROPLET-ASSOCIATED PROTEIN 1 to 3 (LDAP1 to 3) and LDAP INTERACTING PROTEIN (LDIP) that are involved in proper LD formation at the ER (Pyc *et al*., 2017; Pyc *et al*., 2021), as well as PLANT UBX DOMAIN-CONTAINING PROTEIN 10 (PUX10) and CYCLOARTENOL SYNTHASE 1 (CAS1), that play a roles in LD protein degradation (Deruyffelaere *et al*., 2018; Kretzschmar *et al*., 2018) and sterol synthesis (Babiychuk *et al*., 2008), respectively. Interestingly, certain LDAP isoforms are strongly upregulated under different stresses (Winter *et al*., 2007) and their overexpression or disruption indicated they play a role in drought stress resistance (Laibach *et al*., 2015; Kim *et al*., 2016).

Overall, the study of LD proteins and their physiological functions to date has proven crucial to our understanding of LD biology in plants. However, due to the predominant focus of these efforts on seeds, leaves and pollen tubes, sparingly little is known about LDs in roots.

That is, one of the earliest reports of root LDs is their observation in cress (*Lepidium sativum* L.), where LDs are abundantly found in early differentiating statocytes and then decrease in number during differentiation (Hensel, 1986). In leek (*Allium porrum* L.) seedlings, LD number and TAG content were observed to increase in root tissues when their sterol biosynthetic pathway was chemically blocked with fenpropimorph (Hartmann *et al*., 2002). In cotton (*Gossypium sp.*), LDs of root tissue have also been analyzed in seedlings by direct organelle mass spectrometry (MS) and showed a distinct TAG composition enriched in cyclic fatty acids (Horn *et al*., 2011). Among the potential root LD proteins identified, a root-expressed hydroperoxide lyase of *Medicago truncatula* was confirmed to localize to LDs (De Domenico *et al*., 2007). While these studies suggest that root LDs have distinct functions, much of the root LDs’ function in general is still unclear.

In this study, we show that Arabidopsis root LDs act as a sink for acyl chains during heat stress-induced membrane remodeling, and, simultaneously, that LD-stored SEs strongly decrease to serve as a potential source for sterols in membranes, with the highest relative increase being observed in the acylated steryl glycosides (ASGs). Furthermore, we isolated LD-enriched protein fractions from the Arabidopsis double mutant *trigalactosyldiacylglycerol 1-1 sugar dependent 1-4* (*tgd1-1 sdp 1-4*), which has strongly elevated TAG levels (Fan *et al*., 2014), for subsequent proteomic analyses. In our proteomic analysis of the LDs from this mutant, 34 previously described LD proteins were detected. Furthermore, we selected 24 highly-enriched proteins in the LD fraction that have not been reported to be LD localized for further analysis. For 14 of these candidate LD-associated proteins we observed LD localization in transiently transformed *Nicotiana tabacum* pollen tubes and/or *Nicotiana benthamiana* leaf cells. Amongst these LD proteins were two 2,3-oxidosqualene cyclases, suggesting that LDs in roots are hubs for triterpenoid synthesis. Finally, we observed that the isolated root LDs were also enriched in certain triterpenes and triterpene esters, indicating that root LDs can act as a storage site for non-TAG lipids as well.

## MATERIALS AND METHODS

### Plant lines and growth conditions

Lipidomic and proteomic experiments were carried out with Arabidopsis Col-0 and the *tgd1-1 sdp1-4* double mutant line (Fan *et al*., 2014). Seeds of Arabidopsis lines were surface-sterilized with 6% (w/v) sodium hypochlorite and 0.1% (v/v) Triton X-100 and germinated on half-strength Murashige and Skoog (MS; Duchefa Biochemie, Haarlem, The Netherlands) medium (Murashige and Skoog 1962) containing 0.8% (w/v) agar. Plants were grown at a light intensity of 100 µmol s^-1^ m^-2^ for microscopy and 150 µmol s^-1^ m^-2^ for lipidomics, and a temperature of 23°C. The temperature was shifted for 37 °C for heat stress treatment.

For LD enrichment and subsequent proteomic or lipid analysis, high-yield axenic root cultures of *tgd1-1 sdp1-4* were cultivated by adapting an established protocol (Hétu *et al*., 2005). In short, seeds of the oil-rich mutant *tgd1-1 sdp1-4* were surface sterilized, placed on sterile steel grids on top of solid 1/2 MS + 1% sucrose medium, stratified for 72 h, and subsequently grown for 7 d. One-week-old seedlings were transferred to 100 ml Erlenmeyer flasks on the steel grid and supplemented with 10 ml liquid 1/2 MS + 1% sucrose medium. The culture was agitated at 85 rpm for 11 d with regular exchange of the medium every three days. Afterwards, the medium was changed to 15 ml 1/2 MS + 3% sucrose medium and seedlings were grown an additional 11 d before harvest, exchanging the medium every third day.

### Lipidomic sample preparation and measurements

Lipids were extracted from cut-off roots corresponding to ∼ 5 mg dry weight. To inactivate phospholipase activity, samples were initially incubated in boiling water for 20 min. Lipids were sequentially extracted with 1 ml of chloroform:methanol (1:2, v/v), 1 ml of chloroform:methanol (2:1, v/v), and 1 ml of chloroform. For each extraction step, samples were vortexed thoroughly, centrifuged for 10 min at 1500 g and the supernatants collected in a new tube. To the combined supernatant, 0.75 ml of 300 mM NH_4_CH_3_CO_2_ was added, samples were vortexed thoroughly, centrifuged for 5 min at 1500 g, and the lower phase transferred to a new tube. Extracts were evaporated to dryness and dissolved in chloroform:methanol:300 mM ammonium acetate in H_2_O (300:665:35, v/v/v). The dry weight of the remaining residue was determined and the amount of the internal standard adjusted accordingly. Samples were analyzed via direct infusion nanospray MS on an Agilent 6530 Accurate-Mass Q-TOF LC/MS instrument equipped with a ChipCube interface as previously described (Welti *et al*., 2002; Gasulla *et al*., 2013; Gutbrod *et al*., 2021).

Sterols and other triterpenes were extracted from roots corresponding to ∼ 5 mg dry weight. The material was ground with a glass rod after addition of 1000 µl of MTBE:MeOH (3:1) incubated for 1 h at 4°C with shaking. Then, 5 µg of cholestanol standard and 500 µl of ddH_2_O were added, following by vortexing and centrifugation for 10 min at 4°C. The upper organic phase was recovered into a new tube for measurement and the dry weight of the remaining residue was determined for normalization. To measure free sterols and triterpenes, 50 µl were transferred to a GC-vial and evaporated to dryness. Then, the lipids were dissolved in 50 µl hexane and measured by GC-FID, or they were taken up in 15 µl of anhydrous pyridine, derivatized with 15 µl of MSTFA and measured by GC-MS. For total sterol and triterpene measurements, 50 µl of each sample was first evaporated and saponified with 1 ml of 6% methanolic potassium hydroxide at 80°C for 2 h. Lipids were then recovered by adding 500 µl of ddH_2_O and 1 ml hexane 3 times. The hexane extracts were combined and evaporated under a N_2_ stream, before being treated like above for GC-MS measurements.

GC-MS measurements were performed on an 8890 GC system equipped with a HP-5MS UI 30m column (Agilent) coupled to an Agilent 7250 GC/Q-TOF. Helium was used as the carrier gas at a flow rate of 1.1 ml/min. The inlet temperature was set to 250°C. The oven gradient started at 120°C, held for 1 min, then increased at 15°C/min until 300°C and held for 10 min. The ion source and the transfer line temperature were set to 200°C and 280°C, respectively, and 70 eV was used as the electron energy. The mass-to-charge ratio range was recorded from 50 to 500. GC-FID measurements were performed via 8860 GC system equipped with a HP-5MS UI 30m column (Agilent). Helium was used as the carrier gas at a flow rate of 6.5 ml/min, the inlet temperature was set to 250°C and the oven gradient started at 120°C with an increment of 5°C/min up to 300°C.

### Isolation of root LD-enriched fractions and preparations for proteomic analysis

Harvested root material was separated from other plant tissues and drained from the remaining medium by drying between paper towels and applying gentle pressure. The resulting root pads of two separate cultures were pooled into one biological replicate.

For the protein extraction, all processes and materials were kept on ice. Grinding buffer (50 mM Tris-HCl pH 7.5, 10 mM KCl, 0.4 M sucrose, 200 µM proteinase inhibitor PMSF; Carl Roth, Karlsruhe, Germany) and sea sand were added to the root pads which were subsequently ground. To remove cellular debris and sea sand, the homogeneous suspension was centrifuged for 1 min at 100 g. An aliquot was taken from the supernatant and precipitated in 96 % ethanol at –20 °C, representing the total protein fraction. The remaining supernatant was overlaid with washing buffer (50 mM Tris-HCl pH 7.5, 10 mM KCl, 0.2 M sucrose, 200 µM proteinase inhibitor PMSF; Carl Roth, Karlsruhe, Germany) and centrifuged for 35 min at 100,000 g and 4 °C in a swing-out rotor. The floating fat pad was mechanically collected, emulsified in a small volume of washing buffer, and centrifuged in a fixed angle rotor for 35 min at 100,000 g. The floating fat pad was collected again, taken up in 96 % ethanol to remove fat, and stored at –20 °C to precipitate proteins. This fraction was considered as LD fraction.

Precipitated root protein pellets were subjected to defatting by undergoing two washes with 80 % ethanol, followed by drying and an additional wash with 96 % ethanol. The obtained proteins were dissolved in 6 M urea, 5 % SDS (w/v) solution, and their concentrations were determined using the Pierce BCA protein assay kit (Thermo Fisher Scientific, Waltham, MA, USA). 20 µg were subjected to in gel-digest, as previously described (Shevchenko *et al*., 2006; Rappsilber *et al*., 2007).

### LC-MS/MS measurements

An EASY-nLC 1200 system (Thermo Fisher Scientific, Waltham, MA, USA) in conjunction with an Exploris 480 mass spectrometer (Thermo Fisher Scientific, Waltham, MA, USA) was employed for the LC-MS/MS analysis of root-derived peptides. For this purpose, peptides were separated on 20 cm frit-less silica emitters (CoAnn Technologies, Richland, WA, USA) with a 0.75 µm inner diameter, packed in-house with ReproSil-Pur C_18_ AQ 1.9 µm resin (Dr. Maisch, Ammerbuch-Entringen, Germany). The column was maintained at a constant temperature of 50°C. Elution of peptides was carried out over 115 min using a segmented linear gradient from 0 % to 98 % solvent B (solvent A: 0 % ACN, 0.1 % formic acid; solvent B: 80 % ACN, 0.1 % formic acid) at a flow rate of 300 nl min^-1^. The data-dependent acquisition mode was utilized to acquire mass spectra. For full proteome samples, MS^1^ scans were obtained at an Orbitrap resolution of 120,000, covering a scan range of 380-1500 (m/z). The maximum injection time was set to 100 ms, and a normalized AGC target of 300 % was utilized. Precursors with charge states 2-6 were selectively chosen for fragmentation, and up to 20 dependent scans were acquired. Dynamic exclusion was enabled with an exclusion duration of 40 seconds and a mass tolerance of +/-10 ppm. A 1.6 (m/z) isolation window with no offset was established, accompanied by the application of a normalized collision energy of 30. Acquisition of MS^2^ scans was performed at an Orbitrap resolution of 15,000, while maintaining a fixed First Mass (m/z) of 120. The maximum injection time was 22 ms, and the normalized AGC target was set to 50 %.

### Computational processing of MS/MS data

MS/MS raw data were processed in the MaxQuant software (version 1.6.2.17) for feature detection, peptide identification and protein group assembly (Cox & Mann, 2008). Mostly, default settings were used with additional settings as specified in Suppl. Table S1. The TAIR10 protein database (Lamesch *et al*., 2012) was used as reference for identification. Label free quantification was performed to obtain iBAQ and LFQ values for protein abundances (Cox & Mann, 2008; Schwanhausser *et al*., 2011; Cox *et al*., 2014). Further data analysis was done in Perseus 1.6.2.2 (Tyanova *et al*., 2016). Proteomic raw data can be found in the PRIDE database (Vizcaíno *et al*., 2014) under the identifier PXD051152 (https://www.ebi.ac.uk/pride/). All metadata can be found in Suppl. Table S1.

Protein localization was annotated based on the Plant Proteome Database (Sun *et al*., 2009) as of 14th June 2022. LD localization was assigned based on previous studies (Kretzschmar *et al*., 2018; Fernández-Santos *et al*., 2020; Kretzschmar *et al*., 2020; Scholz *et al*., 2024). riBAQ values of proteins were calculated by dividing all individual iBAQ values in one sample through the sum of all iBAQ values in this sample and multiplying by 1000. Only proteins were considered for further analysis that were detected in all 5 replicates of one of the cellular subfractions.

### Molecular cloning and microscopy of *N. tabacum* pollen tubes and Arabidopsis roots

Open reading frames of selected candidate genes were amplified from Arabidopsis floral, root or leaf cDNA and cloned into pLatMCC-GW (Müller *et al*., 2017) via Gateway cloning as described in Müller et al., 2017. All primers used in this study are listed in Table S2. *N. tabacum* growth, pollen transformation, and pollen tube growth were performed as previously described (Müller *et al*., 2017). Images were taken using either a ZEISS LSM780 confocal microscope or a ZEISS LSM 980 with Airyscan 2 confocal microscope (Carl Zeiss Microscopy Germany GmbH, Oberkochen, Germany). This second microscope was also used for monitoring root LDs via the Airyscan 2 function. Detailed settings for all micrographs are described in Table S3. Roots were stained with 1 µg/ml BODIPY 493/503 in water for 5 min prior to microscopy. LD size and number was quantified in Fiji ImageJ (v1.54): In brief, squares of 100 µm x 100 µm of each root micrograph were selected, the LDs within the selected area were recognized using the Threshold (MaxEntropy) function, and the resulting particles after thresholding were quantified with the particle analysis tool.

### Molecular cloning and candidate localization studies *in N. benthamiana* leaves

Open reading frames of selected candidate genes were amplified from cDNA prepared with Maxima Reverse Transcriptase (Thermo Fisher Scientific) according to manufacturer’s instruction using leaf RNA that had been extracted using the Spectrum Plant Total RNA Kit (Merck KGaA, Darmstadt, Germany). Constructs were amplified with the Phusion High-Fidelity DNA Polymerase (Thermo Fisher Scientific, Waltham, MA, USA) following manufacturer’s instructions. Gateway cloning into the plant binary vectors pMDC32-ChC and pMDC32-ChN was carried out by traditional or fast Gateway® cloning as described in (Müller *et al*., 2017). Vector construction of pMDC32-ChC and pMDC32-ChN has been described previously in (Kretzschmar *et al*., 2020) and (Doner *et al*., 2021), respectively. The construct of MmDGAT2 in pMDC32, which was used for co-expression experiments, has been described in (Cai *et al*., 2019). Localization of candidates was analyzed in 28 day-old leaves of *N. benthamiana* that were transiently transformed by infiltration with *Agrobacterium tumefaciens* (strain LBA4404) harboring candidate expression vectors. *N. benthamiana* plant growth, leaf infiltration, and BODIPY 493/503 staining was performed as previously described (Gidda *et al*., 2016; Kretzschmar *et al*., 2020). Leaves micrographs 3 days-post-infiltration were captured as single (0.4 μm) optical z-sections and saved as 512 x 512 pixel images using a Leica SP5 CLSM (Leica Microsystems). Excitations and emission signals for fluorescent proteins and BODIPY were collected sequentially as single optical sections in double-labeling experiments like those described in (Gidda *et al*., 2016), with no detectable crossover observed at the settings used for data collection.

Binary plasmids for putative N-glycan biosynthetic enzymes (Figure S5, At1g78800, At1g16570, At2g40190, At2g47760, At5g38460) were constructed using Gateway Technology (Invitrogen) with the destination vectors pGWB405m (Nakagawa *et al*., 2007; Segami *et al*., 2014) as previously described (Yamaguchi *et al*., 2025). Each full-length genomic coding region without the stop codon was amplified from Arabidopsis Col-0 genomic DNA. Each entry clone was produced by ligating each PCR product into the pENTR plasmid using an In-Fusion Cloning Kit (Takara, Shiga, Japan), and recombined into the destination vector pGWB405m via LR Clonase II (Invitrogen), creating binary vectors for expressing GFP-fused proteins. The binary vectors were transformed independently into *A. tumefaciens* strain GV3101. The transformed cells harboring each plasmid were infiltrated into the leaves of 3-week-old *N. benthamiana* plants. The transformed cells for expressing α-DOX1-RFP were used as LD-localization marker. Two days post-infiltration, epidermal leaf cells were treated with heat stress (50°C, 30 min) for LD induction and observed under a fluorescence microscope (BZ-X800; Keyence, Osaka, Japan). The fluorescent signal of GFP was examined using a GFP filter (excitation, 450–490 nm; emission, 500–550 nm) and that of RFP was examined using a TRITC filter.

### Extraction, derivatization and measurement of triterpenes from isolated LDs

LD-enriched fractions (derived from ∼5-8 g plant material) and total cellular extracts (derived from the same material but only 1 % of the volume of the sample was taken) were obtained as described above for proteomic samples and extracted two times with 1 ml of methanol and two times with 1 ml ethyl acetate. The methanol and ethyl acetate extracts from each fraction were combined and evaporated under N_2_ stream. These dried extracts were then redissolved in hexane. 0.2 µg of cholestanol was added into the total extract fractions and 1 µg to the LD fractions as internal standard. Triterpenes and free sterols in the samples were then extracted, derivatized and measured by GC-MS while membrane glycerolipids and TAGs were determined by direct infusion nanospray MS as described for lipidomic samples above.

### Statistical analysis of data

Statistical analysis of LD numbers (figure 1 and suppl. figure S1) and metabolite data (figures 2,9 and suppl. figure S2) was carried out using the R Statistical Software (v4.2.2; https://www.R-project.org/), while proteomic data (figure 3) was analyzed with the Perseus software platform (Tyanova *et al*., 2016). Lipidomics data and data on LD numbers of heat-stressed vs. control-treated seedlings were analyzed using Welch’s t-test to compare the different treatments. Adjustments for multiple comparisons were made with the ‘p.adjust’-function of the ‘stats’-package in R, applying the Benjamini-Hochberg correction. Calculation of the PCA plot for metabolites in the LD-enriched fraction was done using the ‘prcomp’-function of the ‘stats’-package, setting the parameters ‘center’ and ‘scale.’ both to ‘TRUE’. Volcano plots of proteomics data were calculated from log_2_-transformed and imputed riBAQ values using the ‘Volcano plot’ visualization tool of Perseus. False discovery rate was set to 0.01, otherwise default settings (two-sided t-test, number of randomizations: 250, no grouping in randomizations, S0: 0.1) were used. The resulting values were exported and visualized in Python (v3.10.6; Python Software Foundation, http://www.python.org) using the libraries ‘Matplotlib.Pyplot’ (Hunter, 2007), ‘Pandas’ (McKinney, 2010), and ‘NumPy’ (Harris *et al*., 2020).

**Figure 1:**
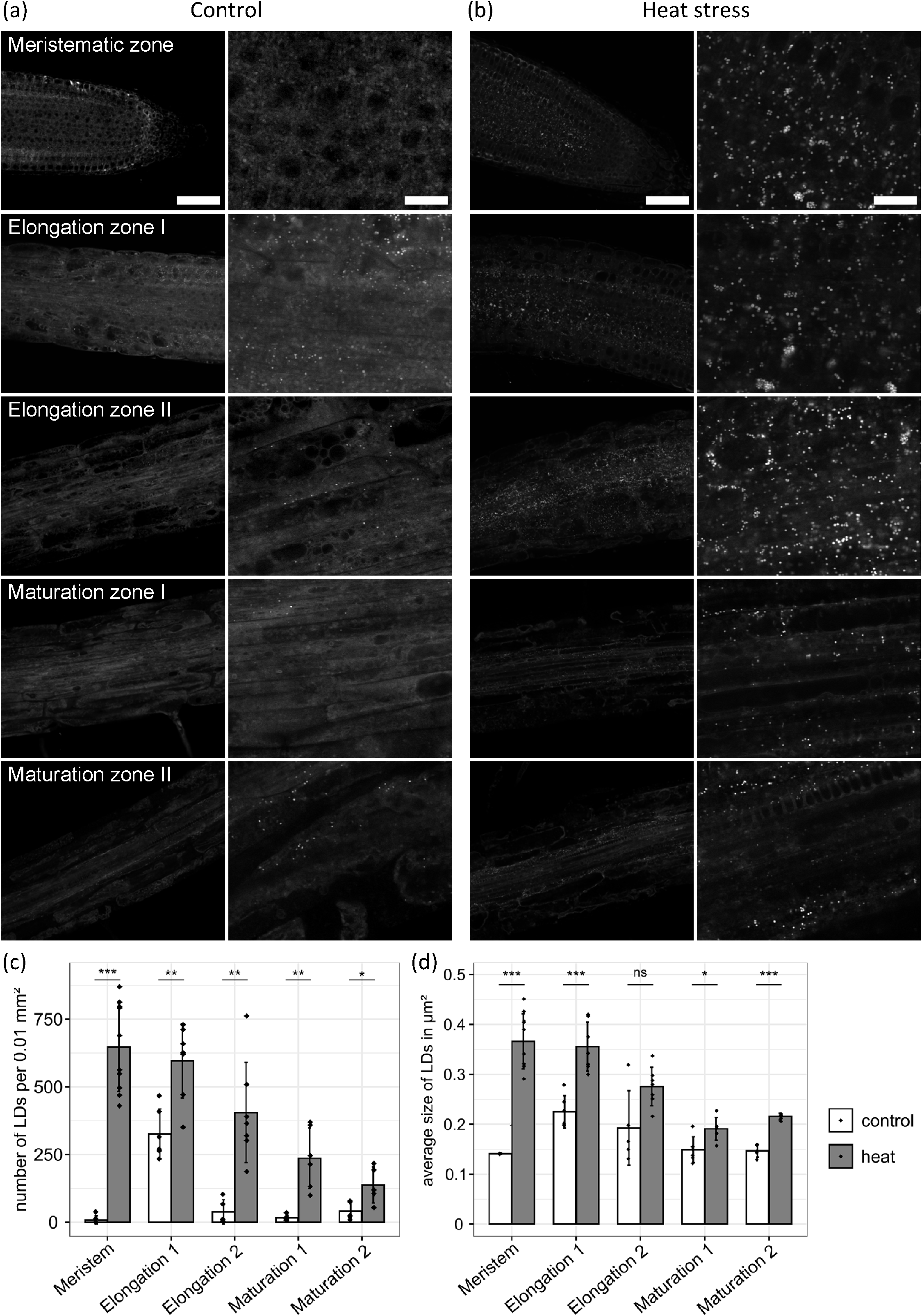
LDs occur in increased number in parts of the root elongation zone and accumulate under heat stress. The roots of 7-day-old Arabidopsis seedlings vertically grown on plates were fixated and stained with BODIPY 493/503. Plants were grown at 23°C (a) and, in the case of heat stress, moved for 24 h to 37°C prior to fixation and analysis (a, b). Median planes of different root zones (see Figure S1) were imaged by CLSM (a, b). For quantitative image analysis, areas up to 100 µm x 100 µm of each root micrograph were selected, and the LDs within the selected areas were quantified in number (c) and size (d). Data was analyzed from n ≥ 5 individual roots, however, in the meristematic zone of control-treated roots, only two roots showed LDs in the micrograph and could be analyzed for LD size in (d). Plots display mean ± standard deviation.

**Figure 2:**
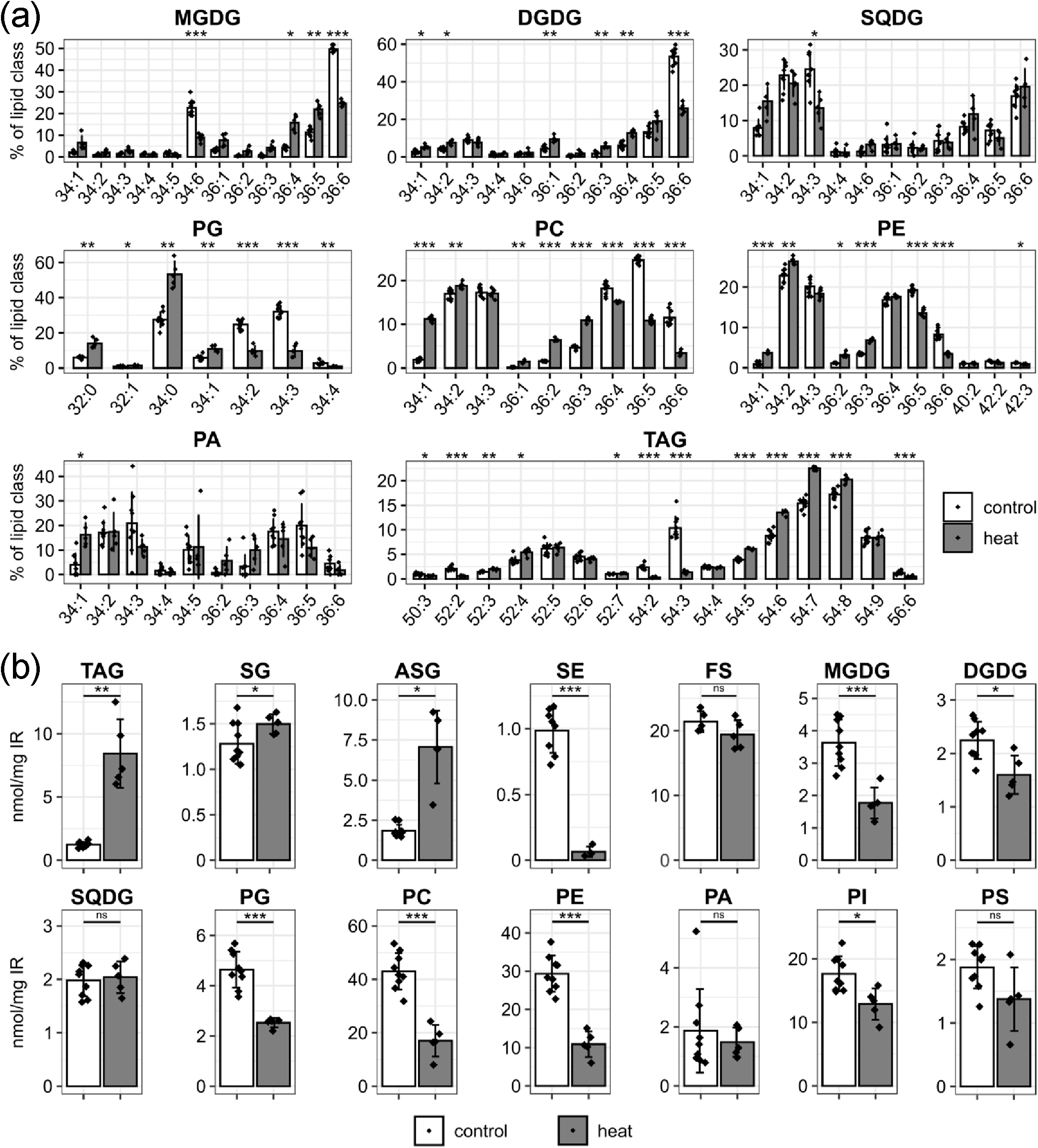
Composition of glycerolipids in control and heat-stressed roots. Lipids were extracted from roots of 12-day-old Arabidopsis seedlings grown vertically on plates. Plants were cultivated at 23°C and, in the case of heat stress, moved for 24 h to 37°C prior to lipid extraction. Lipids were analyzed by ESI-MS/MS. (a) Absolute values for the molar amounts of individual lipid species were determined, and their relative proportion in the respective lipid class was calculated in mol %. (b) Total amounts of lipid classes were determined as sums of all individual lipid species of the respective lipids class. Values are from n = 5-10 biological replicates, and are shown as mean ± standard deviation. Statistical differences were calculated by Welch’s *t*-test using Benjamini-Hochberg correction for multiple comparisons and are represented as follows: *p* > 0.05 “ns”, *p* < 0.05 “*”, *p* < 0.01 “**”, *p* < 0.001 “***”. ASG, acylated steryl glycosides; DGDG, digalactosyldiacylglycerol; FS, free sterols; MGDG, monogalactosyldiacylglycerol; PA, phosphatidic acid; PC, phosphatidylcholine, PE, phosphatidylethanolamine; PG, phosphatidylglycerol; PI, phosphatidylinositol; PS, phosphatidylserine; SE, sterol esters; SG, steryl glycosides; SQDG, sulfoquinovosyldiacylglycerol; TAG, triacylglycerol.

**Figure 3:**
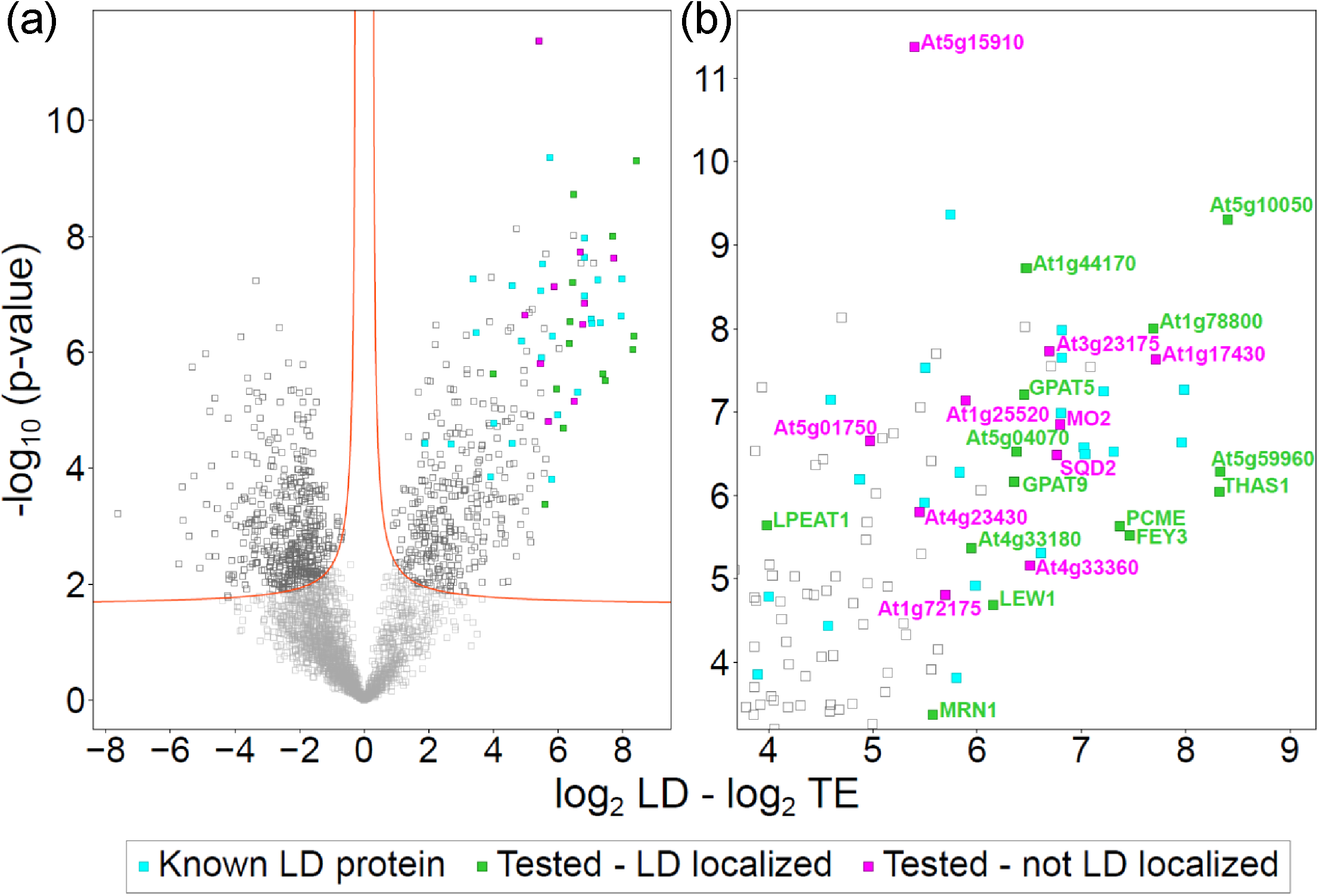
Analysis of protein enrichment in the LD fraction of Arabidopsis roots. The riBAQ dataset was imputed and the values were log_2_ transformed. Then, the difference between the LD-enriched and total protein fractions was calculated for each protein. Additionally, the corresponding *p* values (-log_10_) were determined. (a) A volcano plot was generated based on these values and the upper right corner was enlarged in (b). A false discovery rate (FDR) of 0.01 was used to distinguish between significant and non-significant differences (red lines). Known LD proteins (Table 1) and selected candidate LD proteins (Table 2) are highlighted. FEY3, FOREVER YOUNG 3; GPAT5/9, GLYCEROL-3-PHOSPHATE ACYLTRANSFERASE5/9; LEW1, LEAF WILTING 1; LPEAT1, LYSOPHOSPHATIDYLETHANOLAMINE ACYLTRANSFERASE1; MRN1, MARNERAL SYNTHASE 1, PCME, PRENYLCYSTEINE METHYLESTERASE; THAS1, THALIANOL SYNTHASE 1.

**Table 1:**
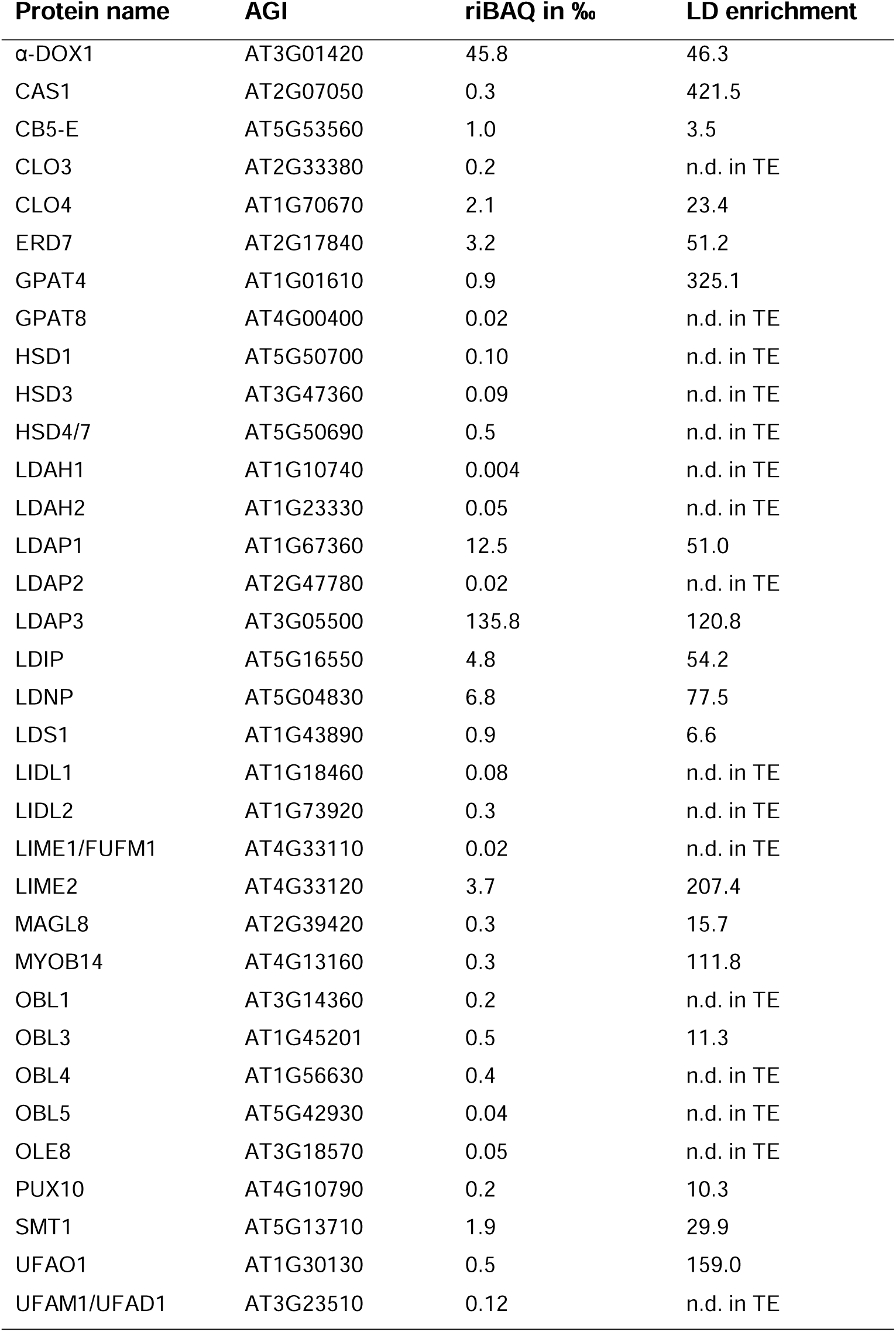
Previously described LD proteins identified in Arabidopsis *tgd1-1 sdp1-4* axenic root cultures. The proteins were derived from LD-enriched (LD) and total (TE) protein fractions. The values provided represent the average riBAQ (n=5) of the LD-enriched fractions. The enrichment of LD proteins was determined by dividing the average LD riBAQ by the average TE riBAQ. n=5.

**Table 2:** Candidate proteins investigated for localization. The proteins were derived from LD-enriched (LD) and total (TE) protein fractions. The values provided represent the average riBAQ (n=5) of the LD-enriched fractions. The enrichment of LD proteins was determined by dividing the average LD riBAQ by the average TE riBAQ. n=5.

| AGI | Acronym | Name | riBAQ LD | Enrichment |
| --- | --- | --- | --- | --- |
| **Detected on LDs through microscopy** |  |  |  |  |
| AT5G60620 | GPAT9 | GLYCEROL-3-PHOSPHATE ACYLTRANSFERASE 9 | 0.54 | 77 |
| AT3G11430 | GPAT5 | GLYCEROL-3-PHOSPHATE sn-2-ACYLTRANSFERASE 5 | 0.32 | n.d. in TE |
| AT1G01610 | GPAT4 | GLYCEROL-3-PHOSPHATE sn-2-ACYLTRANSFERASE 4 | 0.91 | 325 |
| AT1G80950 | LPEAT1 | LYSOPHOSPHATIDYLETHANOLAMINE ACYLTRANSFERASE1 | 0.17 | n.d. in TE |
| AT5G10050 |  | putative short-chain dehydrogenase | 1.23 | 1974 |
| AT5G04070 |  | putative short-chain dehydrogenase | 0.19 | n.d. in TE |
| AT1G44170 |  | putative aldehyde dehydrogenase | 6.55 | 91 |
| AT1G78800 |  | glycosyl transferase family | 1.00 | n.d. in TE |
| AT5G15860 | PCME | prenylcysteine methylesterase | 1.08 | 180 |
| AT4G27760 | FEY3 | FOREVER YOUNG | 0.91 | 132 |
| AT4G33180 |  | putative hydrolase | 0.19 | n.d. in TE |
| AT4G13160 | MYOB14 | MYOSIN BINDING PROTEIN 14 | 0.30 | n.d. in TE |
| AT1G30130 | UFAO1 | UNSATURATED FATTY ACID OXIDASE 1 | 0.51 | n.d. in TE |
| AT5G59960 |  | protein of unknown function | 1.13 | 283 |
| AT1G11755 | LEW1 | LEAF WILTING 1 | 0.26 | 153 |
| AT5G48010 | THAS1 | THALIANOL SYNTHASE 1 | 4.90 | 255 |
| AT5G42600 | MRN1 | marneral synthase 1 | 0.31 | n.d. in TE |
| **Not detected on LDs through microscopy** |  |  |  |  |
| AT4G38540 | MO2 | monooxygenase 2 | 2.97 | 97 |
| AT1G17430 |  | putative hydrolase | 0.43 | n.d. in TE |
| AT5G01220 | SQD2 | SULFOQUINOVOSYLDIACYLGLYCEROL 2 | 3.71 | 102 |
| AT4G23430 |  | putative short-chain dehydrogenase | 5.01 | 40 |
| AT5G15910 |  | putative short-chain dehydrogenase | 3.83 | 42 |
| AT1G72175 |  | putative zinc finger protein | 0.15 | n.d. in TE |
| AT1G25520 | PML4 | PHOTOSYNTHESIS-AFFECTED MUTANT 71 LIKE 4 | 0.20 | n.d. in TE |
| AT3G23175 |  | lesion inducing protein-related | 0.25 | n.d. in TE |
| AT5G01750 |  | protein of unknown function | 2.53 | 29 |
| AT4G33360 | FLDH | farnesol dehydrogenase | 3.51 | 75 |

## RESULTS

### Heat stress in Arabidopsis roots leads to LD proliferation, membrane remodeling, and the accumulation of TAG

Plant LDs have been primarily studied in seeds and also more recently in leaves (Guzha *et al*., 2023). By contrast, LDs in roots are not well explored. We therefore aimed to get an overview of LD abundance in the different developmental zones of the root in Arabidopsis (Figure S1). We imaged BODIPY 493/503-stained LDs of 7-day-old seedlings that were vertically grown on 1/2 MS without sucrose. In most zones of the root, low numbers of LDs were observed, however, in the early elongation zone (adjacent to the meristem), we observed a substantial increase in the number of LDs, reaching an average count of 327 LDs per defined area of 100 x 100 µm² within the single-plane micrograph of the root zone. To investigate potential connections between the number of observed LDs and membrane lipid adaption and turnover in roots, we analyzed LD abundance in heat-stressed roots after incubating the seedlings for 24 h at 37 °C (Figure 1a). Following the heat stress treatment, the number and size of LDs greatly increased throughout the whole root and especially in the meristematic zone (Figure 1b, c).

As these results implicated LDs as a possible sink during heat-induced lipid remodeling, we analyzed the root lipidome of similarly heat-stressed or control seedlings in a targeted manner by ESI-MS/MS and GC-MS (Data S1-S6). While the lipids were measured quantitatively, we first assessed the relative compositions of lipid species within the individual lipid classes, since adaptions here are important for the regulation of membrane fluidity during stress (Figure 2a). The results indicate that 18:3 containing molecular lipid species strongly decreased, while 16:0– and 18:0-containing species relatively increased after heat-stress. These changes are especially pronounced in phosphatidylcholine (PC), where the relative abundance of the 36:5 and 36:6 species dropped from 25 % to 11 % and 12 % to 3 %, respectively. At the same time, for example, 34:1 and 36:2 species increased from 2 % each to 11 % and 6 %, respectively. In TAG, this trend was not observed. Instead, species containing more saturated acyl chains decreased, as the relative proportion of 54:3 and 54:2 dropped from 9 % to 1 % and 2 % to 0.3 %, respectively. For most lipid classes, the average number of double bonds decreased in all membrane lipids. For instance, in phosphatidylcholine from 3.8 to 3.0 or in monogalactosyldiacylglycerol from 5.4 to 4.3 (Figure S2). In TAG, the opposite trend occurred, as the average number of double bonds increased from 5.7 to 6.4. In addition, the absolute amounts of most membrane lipid classes decreased while TAG levels increased 7-fold (Figure 2b). Regarding sterol metabolism, the trend was reversed. Here, the storage form, SEs, was almost completely depleted, while the membrane-localized sterol forms increased, foremost the acylated steryl glycosides (4-fold). Regarding the sterol moiety of the SEs, sitosterol was the most abundant sterol, however, it was also most strongly depleted after heat stress (Data S6).

### Root LD fractions contain numerous strongly-enriched proteins

The remarkable shifts in lipid composition coupled with enhanced LD abundance following heat stress in Arabidopsis roots indicate that LDs can act as both a sink and source for acyl chains and sterols, respectively. Next, we sought to identify additional functions of root LDs by studying their proteome. For LD isolations, high yields of biomass are required. Furthermore, the relatively low number of LDs in the root tissue (Figure 1) needed to be elevated to obtain a floating LD-enriched phase suitable for LD protein isolation. In order to achieve this, we utilized an axenic root culture in high-sucrose media (Hétu *et al*., 2005) of the oil-rich mutant line *tgd1-1 sdp1-4* (Fan *et al*., 2014). The LD-enriched and total cellular fractions were collected from five biological replicates for proteomic analysis. Of the resulting MS data, relative iBAQ values were calculated as ‰ of the summed iBAQ values of all proteins. In total, 4734 different protein groups were identified (Dataset S7), 4318 in the LD fraction and 4134 in the total cellular fraction. In the LD fraction, we detected 34 proteins groups that have been previously reported as LD-associated (Table 1). These annotated proteins were equivalent to 22.4 % of all proteins in the LD fraction but only 0.33 % in the total fraction, implying effective enrichment of LDs by a factor of ca. 70. In order to unravel if any other organelles were systematically co-enriched, we binned the abundance of all proteins with the same assigned subcellular localization (according to the plant proteome database (PPDB; http://ppdb.tc.cornell.edu/; Sun *et al*., 2009). This analysis indicated that besides LD proteins only plastoglobular proteins were enriched at more than an average factor of 4 (Figure S3). All other organelles were either only slightly enriched or depleted.

The most abundant, known LD protein in the LD fraction was LDAP3, with LDAP1 and the LDAP interaction partner LDIP also being detected in comparatively high abundance (Table 1). Interestingly, the dioxygenase α-DOX1 was the second most abundant LD protein, even though in seedlings and leaves it was described as being of much lower abundance (Kretzschmar *et al*., 2020; Scholz *et al*., 2024). In contrast, there was no high-abundant protein from the caleosin or oleosin protein families, whose members dominate the LD proteome of seeds, seedlings and leaves (Brocard *et al*., 2017; Kretzschmar *et al*., 2020; Doner *et al*., 2021; Scholz *et al*., 2024).

Considering our proteomic results, we reasoned that other proteins enriched in the LD fraction could potentially also localize to LDs. To ensure data reliability, we implemented a stringent filter that only considered proteins meeting two criteria: first, they must have been identified with at least two unique peptides; and second, they must have been detected in all five replicates of either the LD or the total cellular extract. In order to identify potential LD proteins, differences of log_2_-transformed riBAQ values between LD and total cellular fractions were calculated. Furthermore, the statistical significance of this difference was calculated through Student’s t-test for each protein (Supplemental Dataset S8). Ultimately, significance and differences between samples for each protein were depicted as a volcano plot (Figure 3). 27 of the proteins (Table 2) that had statistical significance and high enrichment factors were chosen for transient expression experiments in tobacco pollen tubes and/or *N. benthamiana* leaves to assess their subcellular localization by established protocols (Müller *et al*., 2017; Scholz *et al*., 2024). In both transient expression systems, the coding sequence of the protein of interest was appended to a fluorescent tag (mCherry) and potential co-localization to LDs was determined using the neutral-lipid-specific dye BODIPY 493/503. In *N. benthamiana* leaves, the *DIACYLGLYCEROL ACYLTRANSFERASE 2* (*DGAT2*) gene from *Mus musculus* was co-expressed to induce the proliferation of LDs (Cai *et al*., 2019).

### Several LD-enriched proteins localized to LDs and/or the ER

Several putative or characterized enzymes were among the candidate root LD proteins tested for subcellular localization, highlighting that LDs could play an active role in root metabolism. Firstly, we analyzed three candidates with acyl transferase activity that are involved in lipid metabolism (Figure 4). In addition to the GPAT enzymes, GPAT4 and GPAT8, which were previously described to be LD-localized in plants (Fernández-Santos *et al*., 2020), GPAT5 and GPAT9 were also highly enriched in the LD-fraction. In contrast to the other GPATs in Arabidopsis, GPAT9 is a member of a distinct family of plant GPATs (Waschburger *et al*., 2018) that is described as the only GPAT enzyme with a preference to add acyl chains to the *sn1*-not the *sn2*-position of glycerol 3-phosphate (Shockey *et al*., 2016; Singer *et al*., 2016). As shown in Figures 4 and S4, GPAT9 localized almost exclusively to LDs in pollen tubes, while in leaves the localization was only partially on the LD surface. GPAT5 and the previously described GPAT4 (Fernández-Santos *et al*., 2020) displayed a localization to both LDs and the ER in pollen tubes (Figures 4, S4) but GPAT5 showed a ring-like pattern surrounding LDs upon transient expression in *N. benthamiana*. Furthermore, the acyltransferase LYSOPHOSPHATIDYLETHANOLAMINE ACYLTRANSFERASE 1 (LPEAT1) was assayed. LPEAT1 transfers acyl chains to lysophosphatidylethanolamine and to some extent also to lysophosphatidylcholine (Stalberg *et al*., 2009; Jasieniecka-Gazarkiewicz *et al*., 2017). Like GPAT5, LPEAT1 localized not only to LDs but also to the ER in pollen tubes, however, targeted clearly to LDs in leaves (Figure 4). Interestingly, AlphaFold-based structure predictions revealed that each of the respective acytransferases has a broad hydrophobic surface on one side of the protein that could be involved in the interaction with the LD monolayer.

**Figure 4:**
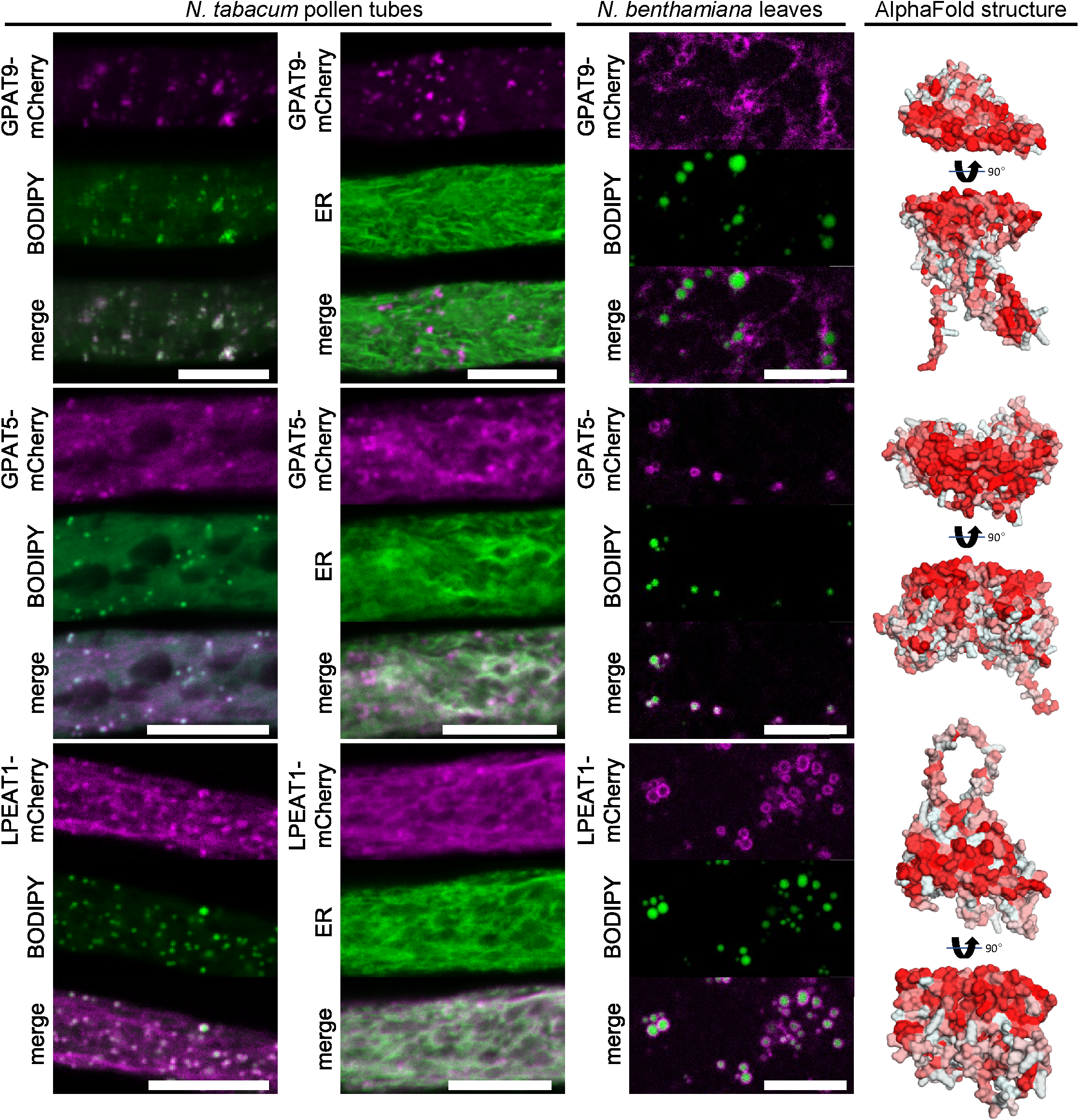
Acyltransferases enriched in the LD fraction of Arabidopsis roots localize to LDs in pollen tubes and leaf cells. mCherry-tagged root LD proteins annotated as acyltransferases were expressed in either *N. tabacum* pollen tubes or *N. benthamiana* leaves. In leaves, the formation of LDs was induced by co-expression with the DIACYLGYLCEROL ACYLTRANSFERASE2 of *Mus muculus* (*Mm*DGAT2). LDs in leaves were subsequently stained with BODIPY 493/503 or the ER was visualized using a co-expressed ER marker (ERD2-CFP). Images are single planes obtained by CLSM. GLYCEROL-3-PHOSPHATE ACYLTRANSFERASE (GPAT9) clearly colocalized with BODIPY-stained LDs in pollen tubes and partially in leaves. GPAT5 and LYSOPHOSPHATIDYLETHANOLAMINE ACYLTRANSFERASE 1 (LPEAT1) localized to LDs and the ER in pollen tubes and to LDs in leaves. Each image is representative at least 6 pollen tubes or 4 leaf areas. Bars, 10 µm. As shown on the right, all three corresponding protein structures, as predicted by AlphaFold2, show a hydrophobic surface on one side of the protein; hydrophobicity indicated by red color.

Further protein candidates that showed localization to LDs at least in *N. benthamiana*, included ALDEHYDE DEHYDROGENASE 4 (ALDH4, At1g44170) and two other proteins annotated by TAIR to contain the NAD(P)-binding Rossmann-fold and, thus, may have putative dehydrogenase functions (Figures 5). The latter two proteins, encoded by *At5g10050* and *At5g04070*, respectively, showed no specific LD-localization when transiently expressed in tobacco pollen tubes, but conversely, displayed ring-like localization around LDs in *N. benthamiana* leaves. ALDH4 showed a similar pattern, although its annular localization around the leaf LDs is more diffuse. AlphaFold predicted protein structures encoded by *At5g10050* and *At5g04070* also both possess a hydrophobic surface not unlike the previously mentioned GPAT5, GPAT9 and LPEAT1 (Figure 4). The predicted structure of ALDH4 is less compact than the other two aforementioned proteins, as one structural domain stretches away from the protein’s center, which contains a hydrophobic region at its distal end.

**Figure 5:**
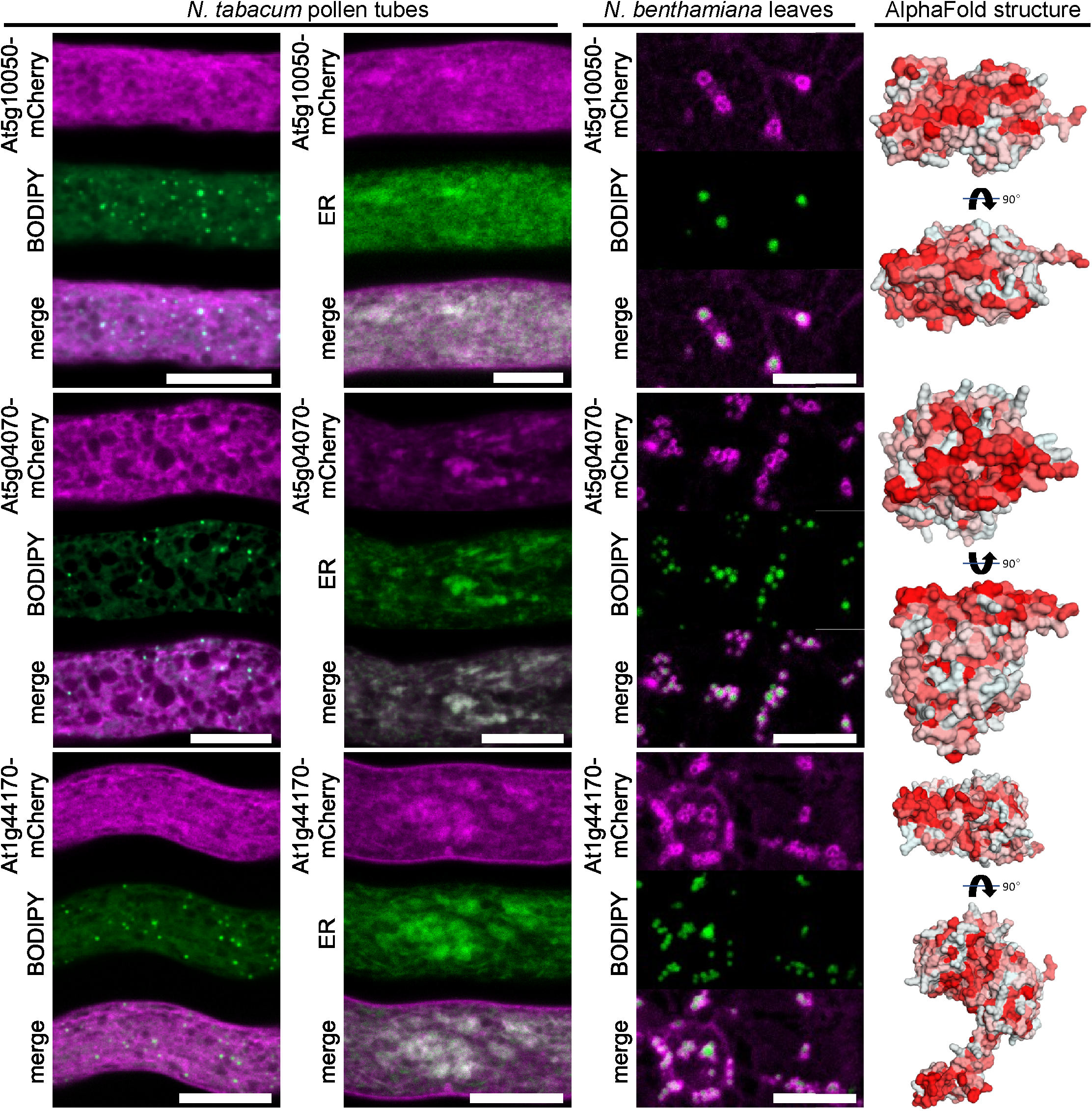
Putative dehydrogenases enriched in the LD fraction of Arabiodpsis roots localize to LDs in leaves. mCherry-tagged proteins were expressed in either *N. tabacum* pollen tubes or *N. benthamiana* leaves. In leaves, the formation of LDs was induced by co-expression with *Mm*DGAT2. LDs were stained with BODIPY 493/503 or the ER was visualized using co-expressed ERD2-CFP. Images are single planes obtained by CLSM. The two putative short-chain dehydrogenases, At5g10050 and At5g04070 and the putative aldehyde dehydrogenase At1g44170 localize to the ER and in the case of At5g10050 and At1g44170 also to the plasma membrane. In leaves, all proteins were localized at LDs. Each image is representative for at least 7 pollen tubes or 4 leaf areas. Bars, 10 µm. As shown on the right, all three corresponding protein structures, as predicted by AlphaFold2, show a hydrophobic surface on one side of the protein, with this surface being at the end of an arm for At1g44170; hydrophobicity indicated by red color.

Additional putative and characterized enzymes that localized to LDs included a glycosyltransferase family protein (At1g78800), the oxidoreductase FOREVER YOUNG (FEY3, At4g27760; Callos *et al*., 1994), a putative hydrolase (At4g33180), and PRENYLCYSTEINE METHYLESTERASE (PCME, At5g15860) that can demethylate isoprenylcysteine methylesters of proteins that have a prenylation as lipid modification at their C-terminal cysteine residues (Deem *et al*., 2006). At1g78800 and PCME localized to LDs, and in the case of At1g78800 also to the plasma membrane of pollen tubes (Figure 6). In addition, when co-expressed with an ER-marker, At1g78800 and PCME also displayed ER co-localization (Figure 6). When expressed in *N. benthamiana*, the fluorescent signal of each individual protein formed clear rings surrounding LDs in leaves. The glycosyltransferase family protein (At1g78800) is an ortholog of yeast ALPHA-1,3/1,6-MANNOSYLTRANSFERASE 2 (Alg2) which is involved in protein N-glycosylation (Gomord *et al*., 2010). We also tested other putative N-glycan biosynthetic enzymes (At1g16570, At2g40190, At2g47760, At5g38460; Gomord *et al*., 2010). The ortholog of yeast Alg1 (i.e., At1g16570) also displayed localization to LDs in *N. benthamiana* leaves (Figure S5), while other enzymes (At2g40190, At2g47760, At5g38460) did not show any localization to LDs (Figure S5).

**Figure 6:**
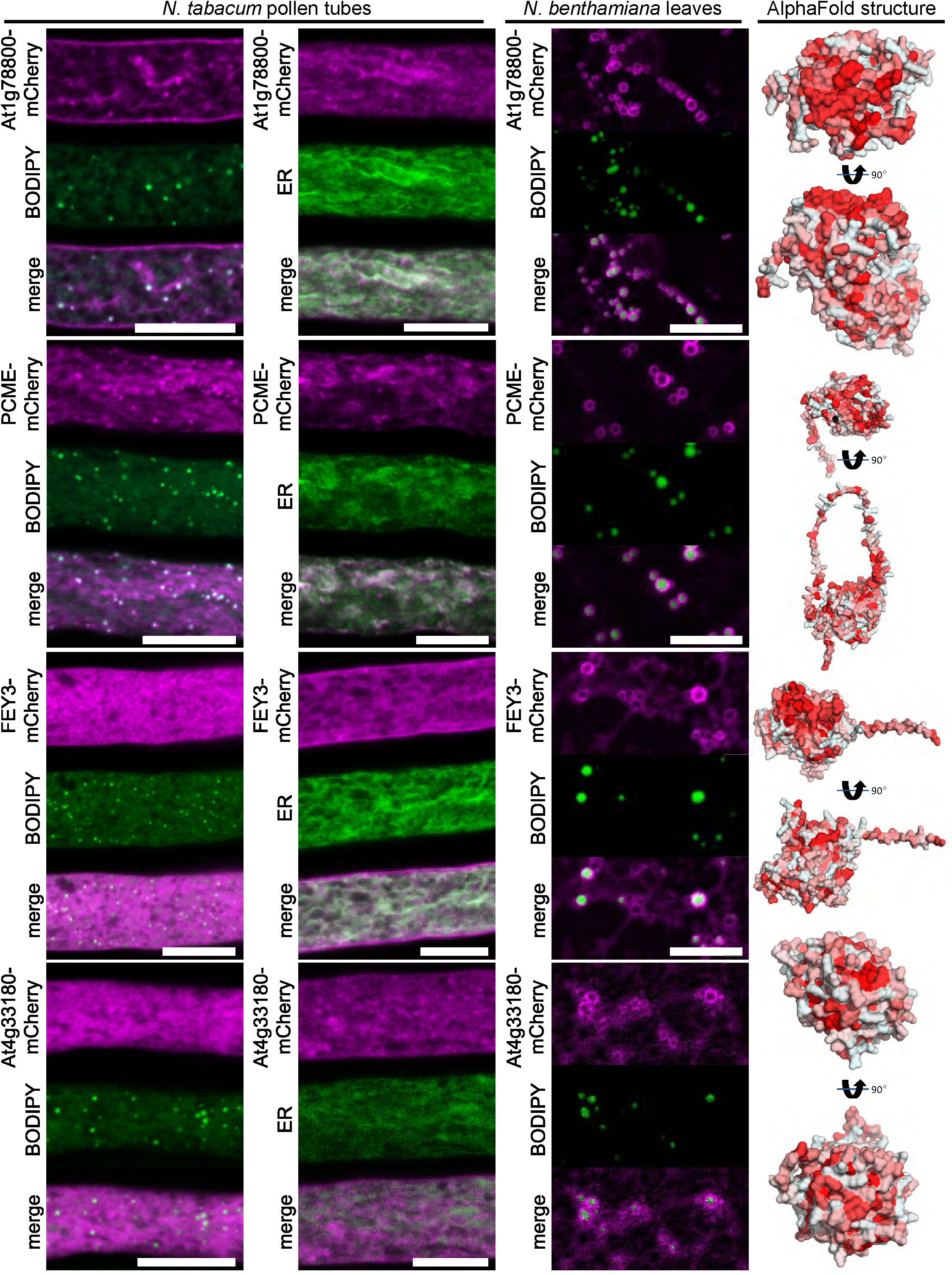
Various enzymes enriched in the LD fraction of Arabiodpsis roots are localized to LDs. mCherry-tagged proteins were expressed in either *N. tabacum* pollen tubes or *N. benthamiana* leaves. In leaves, the formation of LDs was induced by co-expression with *Mm*DGAT2. LDs were stained with BODIPY 493/503 or the ER was visualized using co-expressed ERD2-CFP. Images are single planes obtained by CLSM. In pollen tubes, the putative glycosyl transferase At1g78800, localized to LDs, the ER and the plasma membrane, while the oxidoreductase FOREVER YOUNG3 (FEY3) only localized to the ER and the plasma membrane. The putative hydrolase At4g33180 did not show any colocalization with LDs or the ER but was found in more cloud-like structures and the PRENYLCYSTEINE METHYLESTERASE (PCME) targeted both to LDs and the ER. In leaves, all proteins were localized at LDs. Each image is representative for at least 6 pollen tubes or 4 leaf areas. Bars, 10 µm. As shown on the right, the protein structures of At1g78800 and FEY3, as predicted by AlphaFold2, show a hydrophobic surface on one side of the protein; hydrophobicity indicated by red color.

Localization of FEY3 and a protein of unknown function, At4g33180, expressed in leaves was more diffuse but both accumulated as rings around LDs. However, neither of the two proteins showed an obvious LD localization in pollen tubes (Figure 6). One candidate protein without predicted enzymatic function was MYOSIN BINDING PROTEIN 14 (MYOB14), which has only recently been identified to associate to LDs (Omata *et al*., 2024). Omata et al., also described LD localization for UNSATURATED FATTY ACID OXIDASE 1 (UFAO1), which we also investigated. We could confirm the clear LD association of MYOB14 and UFAO1 in both pollen tubes and leaves (Figure 7). MYOB14 has been proposed as a linker between myosins and LDs, whereby its predicted protein structure suggests that a hydrophobic helix on one side of the protein might interact with LDs, while a putative myosin binding domain could reside in a long adjacent helix. UFAO1 was hypothesized to work together with LIPID DROPLET METHYLTRANSFERASE 1 (LIME1) in furan-containing fatty acid biosynthesis based on homology to a set of proteins studied in the photosynthetic bacterium *Cereibacter sphaeroides* (Lemke *et al*., 2020; Omata *et al*., 2024). Another protein of unclear function, encoded by *At5g59960*, also targeted to LDs in both transient expression systems (Figure 7).

**Figure 7:**
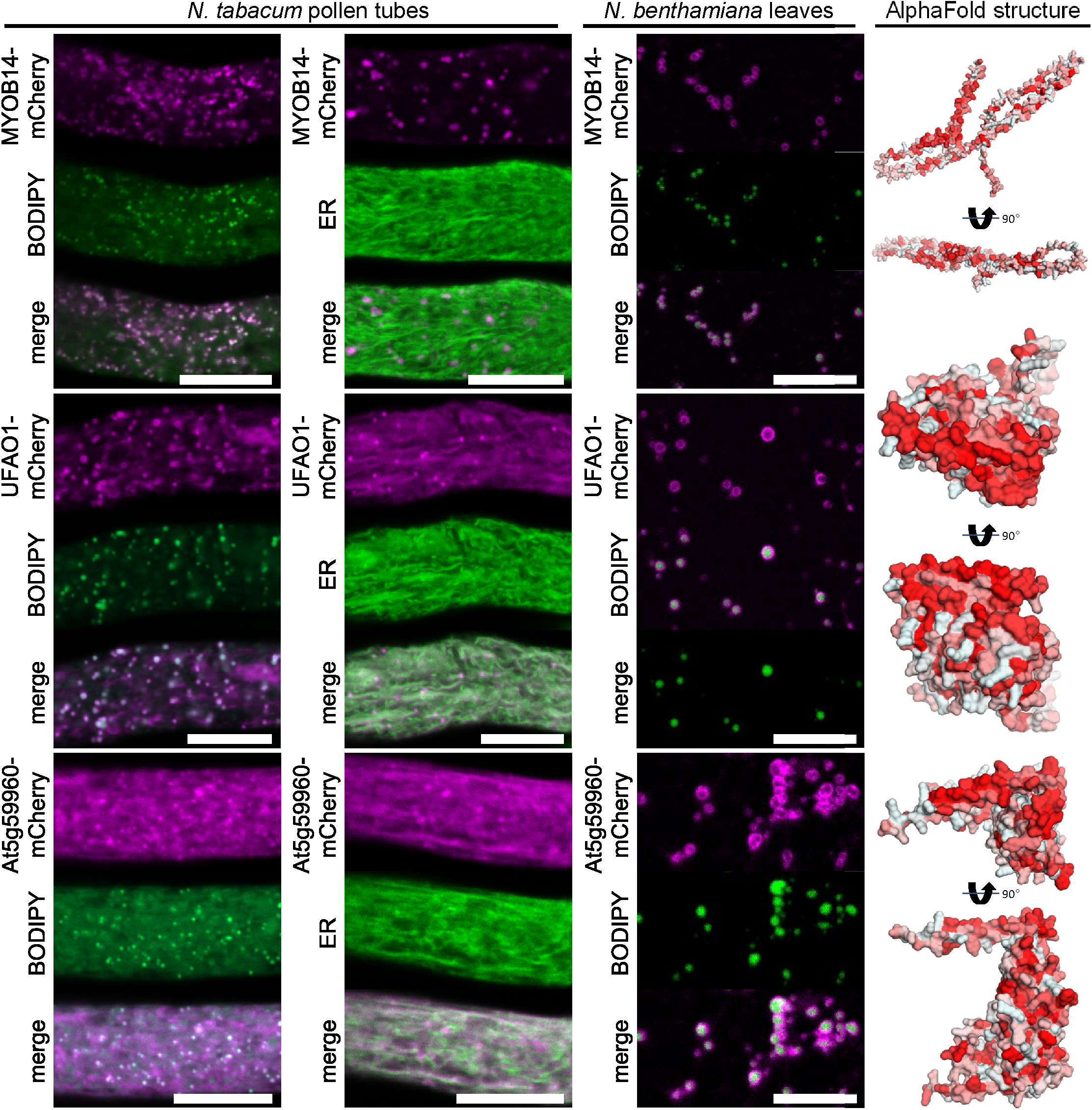
Several proteins of unknown function that are enriched in the LD fraction of Arabidopsis roots are localized to LDs. mCherry-tagged proteins were expressed in either *N. tabacum* pollen tubes or *N. benthamiana* leaves. In leaves, the formation of LDs was induced by co-expression with *Mm*DGAT2. LDs were stained with BODIPY 493/503 or the ER was visualized using co-expressed ERD2-CFP. Images are single planes obtained by CLSM. In pollen tubes, MYOSIN BINDING PROTEIN14 (MYOB14) localized to LDs, while the protein of unknown function At5g59960 and UNSATURATED FATTY ACID OXIDASE1 (UFAO1) localized to LDs and the ER. In leaves, all proteins were localized at LDs. Each image is representative for at least 9 pollen tubes or 4 leaf areas. Bars, 10 µm. As shown on the right, the protein structures of At5g59960 and UFAO1, as predicted by AlphaFold2, show a hydrophobic surface on one side of the protein, while MYOB14 harbors a hydrophobic α-helix; hydrophobicity indicated by red color.

Furthermore, we tested ten additional candidates, including several with described or predicted enzymatic function, that did not localize to LDs (Suppl. Figures S6-S9).

Finally, we investigated three proteins involved in terpenoid metabolism (Figure 8). One of these, the *cis*-prenyltransferase LEAF WILTING 1 (LEW1), which is involved in dolichol biosynthesis (Zhang *et al*., 2008; Kwon *et al*., 2016), localized to the ER in pollen tubes but displayed localization surrounding LDs in *N. benthamiana* leaves. The other two proteins are members of the larger protein family of oxidosqualene cyclases (OSCs), THALIANOL SYNTHASE 1 (THAS1) and MARNERAL SYNTHASE 1 (MRN1).

**Figure 8:**
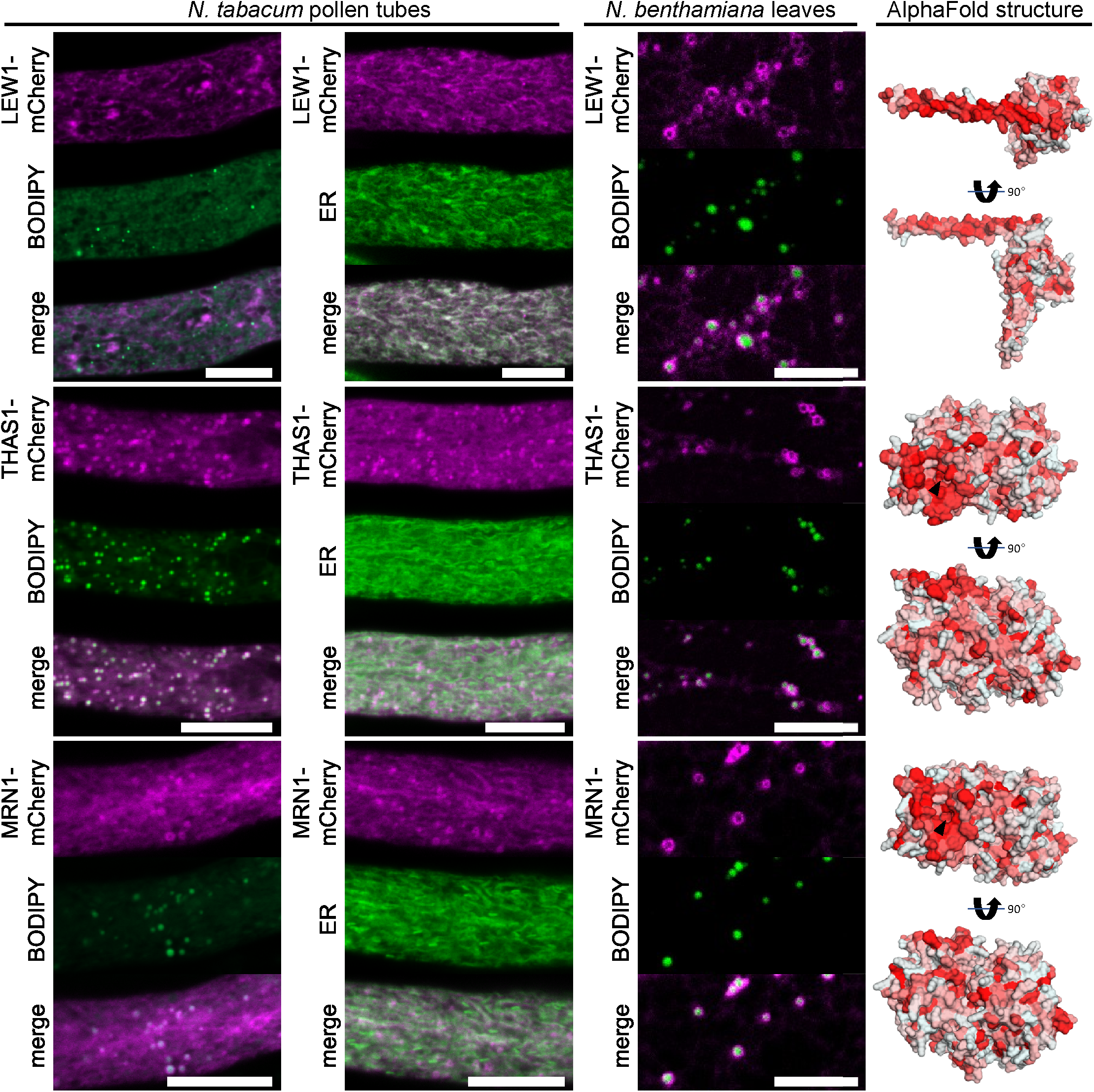
Various enzymes enriched in the LD fraction of Arabidopsis roots that are involved in isoprenoid metabolism are localized to LDs. mCherry-tagged proteins were expressed in either *N. tabacum* pollen tubes or *N. benthamiana* leaves. LDs were stained with BODIPY 493/503 or the ER was visualized by co-expressing ERD2-CFP). Images are single planes obtained by CLSM. THALIANOL SYNTHASE 1 (THAS1) and MARNERAL SYNTHASE1 clearly colocalized with LDs in pollen tubes and in leaves, while the undecaprenyl pyrophosphate synthetase LEAF WILTING 1 (LEW1) targeted the ER in pollen tubes and LDs in leaves. Each image is representative for at least 10 pollen tubes or 4 leaf areas. Bars, 10 µm. As shown on the right, the protein structures of THAS1 and MRN1, as predicted by AlphaFold2, show a hydrophobic surface on one side of the protein (hydrophobicity indicated by red color) with a whole in the middle that might facilitate the uptake of the substrate (black arrow). The structure of LEW1 displays an amphipathic α-helix.

The protein family of OSCs also contains CYCLOARTENOL SYNTHASE, a previously reported LD-localized protein (Kretzschmar *et al*., 2018) that catalyzes the first committed step in phytosterol synthesis. All OSCs share the common substrate (3*S*)-2,3-oxidosqualene, a hydrophobic triterpenoid, but synthesize a strong variety of cyclic triterpenes with 1-5 rings (Hoshino, 2017). THAS1 (Xiang *et al*., 2006) and MRN1 (Xiong *et al*., 2006) have been well explored in regard to their enzymatic function and are both part of gene clusters that harbor genes coding for enzymes that further modify their products (Field & Osbourn, 2008; Huang *et al*., 2019). THAS1 and MRN1 targeted to LDs in both tissues. Interestingly, both THAS1 and MRN1, based on their structural prediction, have a hydrophobic flat face with a hole in the middle that might give access to their substrate.

### Precursors and products of OSCs are enriched in LDs

The localization of THAS1 and MRN1 to root LDs and their reported enzymatic function brought up the question if products and intermediates of the terpenoid metabolism are stored in LDs. To tackle this question, we isolated again root LDs from the *tgd1-1 sdp1-4* mutant but this time analyzed them in respect to their metabolite content using a combination of ESI-MS/MS, and GC-MS. For both membrane and neutral lipids as well as terpenoids, we calculated recovery scores, i.e. how much of the metabolites were recovered in the LD-enriched fractions in comparison to the total fraction in % (Figure 9a).

**Figure 9:**
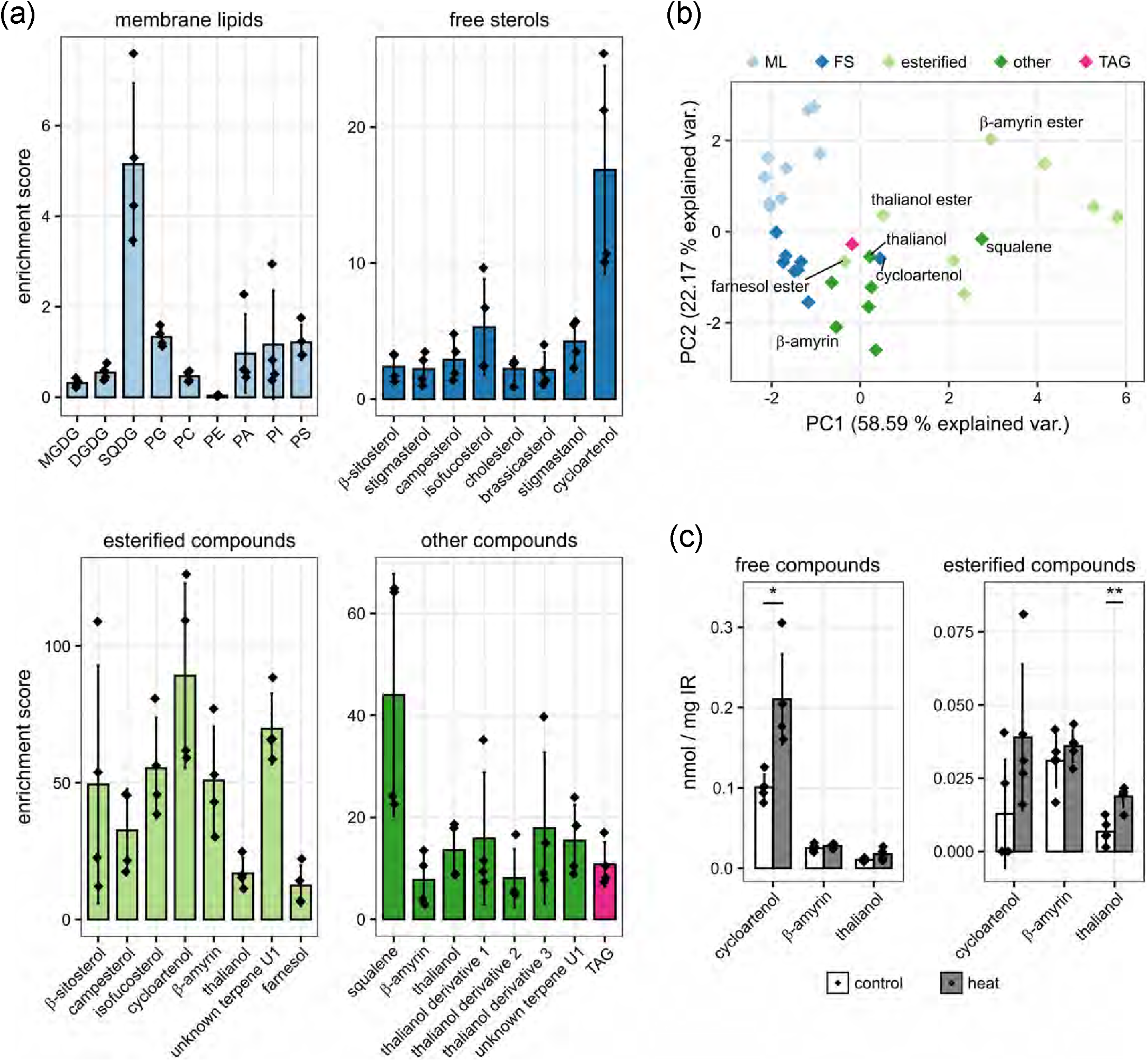
Terpenoids are enriched in LDs isolated from roots of *tgd1-1 sdp1-4* mutant plants. LDs were enriched from roots of the Arabidopsis *tgd1-1 sdp1-4* mutant grown under axenic root cultivation conditions. Aliquots were taken from an initial total cell extract and the final LD-enriched fractions and subsequently analyzed for abundance of various metabolites. Detected compounds were grouped into membrane glycerolipids, TAGs, free sterols, other non-esterified compounds, and esterified compounds. The enrichment scores of individual compounds in the LD-enriched fraction compared to the initial cell extract was calculated (a). The detected quantities of the individual compounds in all samples (normalized to the average of each compound across all samples) were used for a principal component analysis. This analysis supported different enrichment properties of the five compound groups (b). Finally, the three compounds cycloartenol, β-amyrin and thalianol and their esterified forms could also be detected in root lipidomics samples obtained from wildtype seedlings after control and heat-stress treatment (c; same biological samples as initially presented for free sterols in Figure 2). DGDG – digalactosyldiacylglycerol, FS – free sterols, MGDG – monogalactosyldiacylglycerol, ML – membrane lipids, PA – phosphatidic acid, PC – phosphatidylcholine, PE – phosphatidylethanolamine, PI – phosphatidylinositol, PS – phosphatidylserine, SQDG – sulfoquinovosyldiacylglycerol, TAG – triacylglycerol.

The free forms of the major phytosterols, β-sitosterol, stigmasterol and campesterol each had a recovery score of less than 3 % (Supplemental Dataset S10). These values support the notion that the main membrane sterols are present throughout the whole endomembrane system, and do not accumulate in LDs. However, phytosterol recovery is still noticeably higher than membrane phospholipids, which mostly were recovered to less than 1% in the LD-enriched fraction. Remarkably, there were strong differences among the membrane lipids: For example, the recovery of PC was ten times higher than PE indicating a distinct glycerophospholipid composition of root LDs.

Recovery values were drastically higher for esterified sterols, ranging from 33 ± 15 % in campesteryl esters to 89 ± 33 % for cycloartenyl esters. This suggests a clear distinction between free sterols, which are components of cellular membranes throughout the endomembrane system, and steryl esters that are stored in LDs. While higher than membrane lipids, the recovery score of TAG (11 ± 4 %) was notably lower than the steryl esters. Interestingly, certain TAG species were more prevalent in root LDs (Data S11), indicating that select TAG pools are more strongly enriched in the LD fraction than others that might reside in other cellular compartments (e.g., the plastids or the ER).

Precursors of sterols and other triterpenes also appeared to be enriched in LDs, as squalene reached a recovery factor of 44 ± 24 %. Squalene epoxide was also detected in the LD-fraction but levels were too low to be reliably quantified in the total extract. Interestingly, we also found esters of farnesol (at a score of 12.4 ± 7 %) but not free farnesol.

Finally, we were able to identify the two triterpenes β-amyrin and thalianol and their esters. The recovery score of free thalianol was higher than that of most sterols, reaching on average 14 ± 5 %, while β-amyrin was recovered at 8 ± 5 %. The only sterol that reached similar levels to thalianol was cycloartenol (17 ± 8 %), which is also synthesized by an LD-localized enzyme (Kretzschmar *et al*., 2018). In addition, several putative thalianol derivatives had recovery scores ranging from 8 to 18 %. Unlike steryl esters, esters of thalianol (14 ± 5 %) were recovered from LDs with similar efficiency as free thalianol, implicating that both molecules are likely similarly associated with LDs. Recovery of β-amyrin esters (51 ± 20 %) was remarkably greater than free β-amyrin. A principal component analysis (PCA) of the values across samples also showed that the different metabolite classes exhibit similar patterns of enrichment in LDs (Figure 9b).

As the LDs analyzed for their composition were derived from axenic roots grown under high sucrose levels, we aimed to additionally identify triterpenes in Arabidopsis roots grown under more standard conditions, i.e. on agar plates. For this, we reanalyzed the GC-MS data on lipid extracts previously obtained for free sterol analysis (Figure 2, Data S5). Indeed, we were able to identify both the esterified and free forms of thalianol and amyrin (Figure 9c, Data S5), albeit at much lower levels than for phytosterols (Figure 2). In conclusion, these proteomic and metabolomic data support the notion that OSCs, their precursors and direct products (and their esters) are enriched on and in LDs.

## DISCUSSION

### LDs act simultaneously as a sink and source during membrane remodeling

Our investigation of root LD composition identified compounds known to accumulate within LDs of other tissues, such as TAGs and SEs, but also distinct proteins and metabolites that hint to them having additional, previously unknown functions.

Based on our initial observations, the early elongation zone of Arabidopsis roots grown under control conditions accumulate LDs to a much higher extent than other root zones (Figure S1). The reason for this local maximum of LDs is unclear, however, it is tempting to speculate that they accumulate here prior to rapid cell elongation, which demands resources for membrane lipid synthesis. Similarly, it has been suggested that LDs in pollen tubes help deliver precursors of membrane phospholipids to the apical membrane which is constantly extended (Ischebeck, 2016). LD numbers in all zones of the Arabidopsis root increase after plants have been subjected to one day of heat stress (Figure 1). Concomitantly, we observed an increase in the total amount of TAGs in roots (Figure 2), comparable to previous reports from Arabidopsis seedlings and leaves (Mueller *et al*., 2015; Shiva *et al*., 2020; Scholz *et al*., 2024), and tobacco pollen tubes (Krawczyk *et al*., 2022a). As such, processes in roots are probably similar to the ones proposed in leaves, albeit with less contributions of plastidial lipids: LDs and its core component TAG serve as sink to sequester acyl chains from degraded membrane lipids, as the membrane is remodeled to adapt to the change in temperature. Supporting this hypothesis, the average number of double bonds in root TAGs increased after heat stress, while the average number of double bonds in membrane lipids decreased, as polyunsaturated acyl chains are channeled from membrane lipids into TAGs to decrease membrane fluidity at higher temperatures (Figure S2).

Interestingly, SEs that occur at similar levels as TAG in unstressed roots decrease during heat stress (Figure 2). This suggests that SEs act as a source for the increase in acylated sterol glycosides (Suppl. Dataset S6). While the cellular function of acylated sterol glycosides is unclear, one could speculate that they stabilize membranes or membrane domains under stress. In conclusion, LDs might simultaneously act as a sink and source to sustain membrane lipid homeostasis under stress.

### Root LDs are distinct from LDs of other organs

When comparing the root LD proteome (Table 1) to LD proteomes of other tissues, (Brocard *et al*., 2017; Kretzschmar *et al*., 2018; Kretzschmar *et al*., 2020; Scholz *et al*., 2024) it appears most similar to leaves, primarily because no oleosins are found and LDAPs (Gidda *et al*., 2016) are the main LD proteins. Nevertheless, there are some striking differences also between LDs of Arabidopsis leaves and roots. Firstly, in leaf LDs, CALEOSIN3 is the most abundant LD protein (Scholz *et al*., 2024), while in roots, the two detected proteins of the caleosin protein family comprised only minor amounts of the LD protein fraction (Table 1). Conversely, α-DOX1 is the second most abundant protein in root LDs, while it is only present in small amounts in leaves (Scholz *et al*., 2024). There, α-DOX1 and CLO3 have been described to act in concert in the oxidation of α-linolenic acid to 2-hydroxy-octadecatrienoic acid (Shimada *et al*., 2014; Shimada *et al*., 2015). Notably, in tomato, the expression of an α*-DOX1* gene was reported to be responsive to salt stress and wounding (Tirajoh *et al*., 2005), thus it cannot be ruled out that the conditions of our root cultures similarly induced α*-DOX1* expression. Nevertheless, even if the hyper accumulation of α-DOX1 in root LDs is stress-mediated, the independence of its accumulation from high protein amounts of caleosins is quite striking and in contrast to leaf LDs, where both α-DOX1 and CLO3 are upregulated after different stresses (Scholz et al., 2024). Hence, putative reaction products of α-DOX1 might be processed differently at root LDs compared to leaf LDs.

Further differences in metabolic reactions at LDs are implied by a number of root LD proteins that were not detected in leaves. These include steroleosins, for which several isoforms were detected. HSD4 and/or 7 were most abundant. These two proteins are identical on the protein level and therefore not distinguishable in proteomic datasets. Regarding HSD4/7, it is interesting to note that these particular steroleosins have not been reported in other plant tissues so far, even though other members of the protein family have been reported in seeds and seedlings (Baud *et al*., 2009; Kretzschmar *et al*., 2020). For the Arabidopsis steroleosin HSD1, hydroxysteroid dehydrogenase activity on mammalian sterols has been reported (d’Andrea *et al*., 2007), which led to speculation that steroleosins could have a role in the conversion of different brassinosteroids thereby regulating their biological activity (Chapman *et al*., 2012). However, no endogenous substrates of any steroleosin have been identified so far, thus a potential role of HSD4/7 in the regulation of root brassinosteroid activity is highly speculative.

Other proteins like GPAT9 and LPEAT1 were identified as LD associated in roots in this study but have clearly important functions in all plant cells (Shockey *et al*., 2016; Jasieniecka-Gazarkiewicz *et al*., 2017). These proteins and others localize not only to LDs but also the ER (Figures 4-8), raising the question why this dual targeting is observed. One reason could be that due to overexpression effects, the binding sites on the LDs get saturated leading to ER targeting instead. Furthermore, proteins might bind the ER first and then move over to the LDs as proposed for oleosin (Beaudoin & Napier, 2002) and various mammalian LD proteins (Kory *et al*., 2016; Song *et al*., 2022), and are imaged while *en route*. Another possibility is that proteins shuttle between LDs and the ER as a mechanism to regulate their activity. For example, GPAT9 might reside inactively at LDs before moving to the ER, where it more likely finds its substrate acyl-CoA (Bates, 2016).

### Arabidopsis root LDs are hubs for triterpene synthesis and storage

Apart from enzymes involved in glycerolipid synthesis, we found enzymes of terpenoid metabolism at root LD, including the 2,3-oxidosqualene cyclases MRN1 and THAS1 (Table 2, Figure 8). These proteins share strong sequence similarities with the 2,3-oxidosqualene cyclase cycloartenol synthase which catalyzes the first committed step in phytosterol biosynthesis and is also localized at LDs (Table 1; Kretzschmar *et al*., 2018; Kretzschmar *et al*., 2020; Scholz *et al*., 2024).

Both *MRN1* and *THAS1* are part of gene clusters, wherein adjacent genes encode enzymes that modify the initial reaction product of MRN1 or THAS1 (Field & Osbourn, 2008; Huang *et al*., 2019). *THAS1* expression in particular appears to be root-specific and the metabolites derived from thalianol have been reported to modify the Arabidopsis root microbiome (Huang *et al*., 2019). The substrate of both enzymes, 2,3-oxidosqualene and its precursor squalene, are highly hydrophobic and appear to be enriched in the core of root LDs (Figure 9). The discrete hydrophobic surfaces of these individual enzymes may enable LD binding and as a result, allow these enzymes to access metabolites stored within the LD cores through a hole in this surface (Figure 8). Interestingly, their products, which are amphipathic due to a hydroxy group, are also enriched in LD fractions (Figure 9). This was also found for cycloartenol indicating that the products of 2,3-oxidosqualene cyclases reside in the LDs for some time before moving to the ER where the downstream enzymes are localized (Figure S10). Alternatively, the products of 2,3-oxidosqualene cyclases could get esterified directly at the LDs, as esters of triterpenes were also enriched in LDs. In conclusion, root LDs appear as prime hubs for triterpene synthesis and storage in Arabidopsis. In other plants they might have this major function also in other tissues, as for example rosemary leaves, birch bark and olive fruit store high amounts of triterpenes (Jäger *et al*., 2009).

## Supporting Information

Fig. S1. Lipid droplets are enriched in parts of the elongation zone

Fig. S2. The average number of double bonds decreases in most lipid classes under heat stress.

Fig. S3. Enrichment of different organellar proteomes in the LD-enriched fraction.

Fig. S4. Subcellular localization of GLYCEROL-3-PHOSPHATE ACYLTRANSFERASE 4 (GPAT4).

Fig. S5. Subcellular localization of N-glycan biosynthetic enzymes.

Fig. S6. Subcellular localization of further candidates.

Fig. S7. Subcellular localization of putative dehydrogenases.

Fig. S8. Subcellular localization of further candidate proteins with unknown function.

Fig. S9. Subcellular localization of further candidate proteins with unknown function.

Fig. S10. Subcellular localization of enzymes acting downstream of THAS1.

Table S1. Metadata for proteomic analysis

Table S2. List of primers used in this study

Table S3. Microscopy settings pollen tubes

Dataset S1: Membrane lipids – absolute levels.

Dataset S2: Membrane lipids – relative composition of species per class.

Dataset S3: Triacylglycerols – absolute levels.

Dataset S4: Triacylglycerols – relative composition of species.

Dataset S5: Free sterols, and triterpenoids and their esters – absolute levels and relative composition.

Dataset S6: Sterol derivatives.

Dataset S7: Proteins found in Arabidopsis roots of the *tgd1-1 sdp1-4* mutant – normalized riBAQ and rLFQ values.

Dataset S8: Comparison of proteins in LD-enriched fractions to total protein fractions isolated from Arabidopsis *tgd1-1 sdp1-4* roots. Log_2_ transformed and imputed data.

Dataset S9: Terpene and sterol enrichment in LDs – raw values.

Dataset S10: Terpene and sterol enrichment in LDs – processed data.

Dataset S11: Triacylglycerol (TAG) enrichment in LDs – processed data.

Dataset S12: Membrane lipid enrichment in LDs – processed data.

## ACKNOWLEDGMENTS

We thank Ivo Feussner (University of Göttingen) for hosting and supporting the Ischebeck lab during parts of this work, and Jan-Ole Niemeier and Markus Schwarzländer for support with microscopy. We would like to thank Paulina Heinkow for technical assistance, especially in maintaining the LC-MS/MS instruments at the Mass Spectrometry-based Proteomics Unit Biology of Plants (MSPUB). Shoji Mano (National Institute for Basic Biology), Shoji Segami (National Institute for Basic Biology), Tsuyoshi Nakagawa (Shimane University), Sumie Ishiguro (Nagoya University) for donating vectors.

## AUTHOR CONTRIBUTIONS

P.S., P.W.N., T.L.S., I.F., P.D., R.T.M. and T.I. designed the research; P.S., J.D., A.C.C., P.W.N., A.C.V., M.S.S.L., S.S., L.H., F.D., K.F.B., L.P., E.L., Y. I. and K.G. performed research; P.S., P.W.N., A.C.V., M.S.S.L., M.B., T.L.S., J.E., I.F., K.G., P.D., R.T.M. and T.I. analyzed data; P.S., T.L.S. and T.I. wrote the paper with the help of all authors. P.S., J.D. and A.C.C. contributed equally.

## DATA AVAILABILITY

Data are available in the article supporting material and the proteomic raw data under the identifier PXD051152.

