## Supplementary material for "Arabidopsis root lipid droplets are hubs for membrane homeostasis under heat stress, and triterpenoid synthesis and storage": Table S

| **Supplemental Table S1: Metadata file for LC-MS/MS data processing with MaxQuant.** Proteome of *Arabidopsis thaliana tgd1-1 sdp1-4* roots, total cellular extracts and lipid droplet-enriched fractions | |
|---|---|
| **1. General features** | |
| Responsible persons | Prof. Till Ischebeck[1], Dr. Jürgen Eirich[2], Prof. Dr. Iris Finkemeier[2]<br>[1]Institute of Plant Biology and Biotechnology (IBBP), University of Münster, Green Biotechnology, Münster 48143, Germany<br>[2]Institute of Plant Biology and Biotechnology (IBBP), University of Münster, Plant Physiology,, Münster 48143, Germany |
| Instrument manufacturer, model | Thermo Fisher Scientific, Orbitrap Exploris 480 |
| Experimental Design | Analysis of total cellular and lipid droplet-enriched fractions of Arabidopsis *tgd1-1 sdp1-4* roots grown as axenic root culture approach in high-sucrose media |
| Groups | Total protein, lipid droplet-enriched fraction |
| Biological and technical replicates | Biological replicates: 5 for each subcellular fraction<br>PS163, 165, … total cellular fractions<br>PS162, 164, … lipid droplet enriched fractions |
| Sample amount | 10 |
| **2. Electrospray Ionisation (ESI)** | |
| Supply type (static or fed) | fed |
| Interface manufacturer | Thermo Fisher Scientific |
| Sprayer type | Nanospray Flex Ion Source |
| **3.1 Post source component – Analyser** | |
| | Orbitrap Exploris 480: Orbitrap analyser |
| **3.2 Post source component – Activation/dissociation** | |
| Instrument component where the activation/dissociation occurs | Orbitrap Exploris 480: HCD cell |
| Gas type | Orbitrap Exploris 480: Nitrogen |
| Activation/dissociation type | Orbitrap Exploris 480: HCD |
| **4.1 Spectrum and peak list generation and annotation – Data acquisition** | |
| Software name and version | Xcalibur 4.0 |
| Acquisition parameters | Data-dependent Top20 |
| Software name and version | MaxQuant 1.6.2.17 |
| **4.2 Spectrum and peak list generation and annotation – Resulting data** | |
| Location of source and processed files | The mass spectrometry proteomics data will be deposited to the ProteomeXchange Consortium via the PRIDE partner repository |
| **5. Description of the software and methods applied in the quantitative analysis** | |
| Quantification software | MaxQuant 1.6.2.17 |
| Description of the selection and/or | Upload of all .raw files into the software. Grouping of technical replicates as one Experiment (“set experiment”). |

| | |
|---|---|
| matching method of features, together with the description of the method of the primary extracted quantification values determination for each feature and/or peptide | Group-specific parameters:<br>1) Type: default<br>2) Digestion: default<br>3) Modifications: default<br>4) Label-free quantification: LFQ, default<br>5) Instrument: intensity determination: total sum, rest default<br>6) First search: default<br>7) Misc: default<br>Global parameters<br>1) Sequences: updated TAIR10 peptides from 14.12.2010, rest default<br>2) Identification: PSM FDR=0.01, protein FDR=0.01, Match between runs ✓, rest default<br>3) Protein quantification: default<br>4) Label free quantification: iBAQ ✓, rest default<br>5) Tables: default<br>6) Folder locations: default<br>7) MS/MS analyzer: FTMS recalibration ✓, rest default<br>8) Advanced: default |
| Confidence filter of features or peptides prior to quantification | Global parameters: identification: PSM FDR=0.01, protein FDR=0.01, rest default |
| Normalisation | All values were divided by the total iBAQ intensities or total LFQ intensities in one sample and multiplied by 1000. |

| AGI | Acronym | Name | 5' end | 3' end |
|---|---|---|---|---|
| AT5G60620 | GPAT9 | glycerol-3-phosphate acyltransferase 9 | GGGG ACAAGTTTGTACAAAAAAGCAGGCT C ATGAGCAGTACGGCAGGG | GGGG ACCACTTTGTACAAGAAAGCTGGGT CCTTCTCTTCCAATCTAGCCAGGA |
| AT3G11430 | GPAT5 | GLYCEROL-3-PHOSPHATE sn-2-ACYLTRANSFERASE 5 | GGGG ACAAGTTTGTACAAAAAAGCAGGCT C ATGGTTATGGAGCAAGCTGGAAC | GGGG ACCACTTTGTACAAGAAAGCTGGGT C ATGGAGACAAGGCTCGAAAGTG |
| AT1G01610 | GPAT4 | GLYCEROL-3-PHOSPHATE sn-2-ACYLTRANSFERASE 4 | GGGG ACAAGTTTGTACAAAAAAGCAGGCT C ATGTCTCCGGCGAAGAAGA | GGGG ACCACTTTGTACAAGAAAGCTGGGT C CTCCATGGACTTGGTCTTATTGAT |
| AT1G80950 | LPEAT1 | LYSOPHOSPHATIDYLETHANOLAMINE ACYLTRANSFERASE1 | GGGG ACAAGTTTGTACAAAAAAGCAGGCT C ATGGAATCAGAGCTCAAAGATTTGAA | GGGG ACCACTTTGTACAAGAAAGCTGGGT C TTCTTCTTTCTGATGGAAATCACGG |
| AT5G10050 | | putative short-chain dehydrogenase | GGGG ACAAGTTTGTACAAAAAAGCAGGCT C ATGGAGAGTGGCGATGAGAGTC | GGGG ACCACTTTGTACAAGAAAGCTGGGT C CTTCTTCATTAAAACCTGCTTCTGG |
| AT5G04070 | | putative short-chain dehydrogenase | GGGG ACAAGTTTGTACAAAAAAGCAGGCT C ATGGAGAATTTGAAGGAGGCT | GGGG ACCACTTTGTACAAGAAAGCTGGGT C AGTGTGAGTTTGCTGCAATT |
| AT1G44170 | | putative aldehyde dehydrogenase | GGGG ACAAGTTTGTACAAAAAAGCAGGCT C ATGGCTGCGAAGAAGGTTTTTG | GGGG ACCACTTTGTACAAGAAAGCTGGGT C AGCTAAACCGAGAAGGACTTTG |
| AT1G78800 | | glycosyl transferase family (gateway) | GGGG ACAAGTTTGTACAAAAAAGCAGGCT C ATGGCGAAAAAAGAAGGTTCAAAG | GGGG ACCACTTTGTACAAGAAAGCTGGGT C ATCTTCTTTAGGACTTGATACGAC |
| AT1G16570 | | putative N-glycan biosynthetic enzyme | GCAGGCTCCGCGGCCATGGGGAAAAGAGGAAGGGCT | AGCTGGGTCGGCGCGTGAATCTGCAATTTGAGACAC |
| AT1G78800 | | glycosyl transferase family | GCAGGCTCCGCGGCCATGGCGAAAAAAGAAGGTTCA | AGCTGGGTCGGCGCGATCTTCTTTAGGACTTGATAC |
| AT2G47760 | | putative N-glycan biosynthetic enzyme | GCAGGCTCCGCGGCCATGGCGGGCGCCTCATCACCG | AGCTGGGTCGGCGCGTGCTTTTTTGTGTATTTGGGA |
| AT5G38460 | | putative N-glycan biosynthetic enzyme | GCAGGCTCCGCGGCCATGCCGAAGAAGAAGCCGGCG | AGCTGGGTCGGCGCGGATTTGCTTCTTTTCTTTATC |
| AT2G40190 | | putative N-glycan biosynthetic enzyme | GCAGGCTCCGCGGCCATGGCGATCTACTTCATTCTC | AGCTGGGTCGGCGCGTTTAAGAGGACCTGTGAAAAT |
| AT5G15860 | PCME | prenylcysteine methylesterase | GGGG ACAAGTTTGTACAAAAAAGCAGGCT C ATGCATTCGCCTCTTCAGACTC | GGGG ACCACTTTGTACAAGAAAGCTGGGT C GAAAGGGCTAATCTCACGAGCC |
| AT4G27760 | FEY3 | FOREVER YOUNG | GGGG ACAAGTTTGTACAAAAAAGCAGGCT C ATGAGTGACGAAACGACGTCATC | GGGG ACCACTTTGTACAAGAAAGCTGGGT C TTCGTGTTGTGCTCCATACCG |
| AT4G33180 | | putative hydrolase | GGGG ACAAGTTTGTACAAAAAAGCAGGCT C ATGTCGCTGTCGTCTTACCTC | GGGG ACCACTTTGTACAAGAAAGCTGGGT C AATATTGTTGAACTCTTGAGCACATTC |
| AT4G13160 | MYOB14 | MYOSIN BINDING PROTEIN 14 | GGGG ACAAGTTTGTACAAAAAAGCAGGCT C ATGGACTACCAAGAAAGTTATAGATTGAC | GGGG ACCACTTTGTACAAGAAAGCTGGGT C TGGGAGATGTGTTGAAGATGAAGT |
| AT1G30130 | UFAO1 | UNSATURATED FATTY ACID OXIDASE 1 | GGGG ACAAGTTTGTACAAAAAAGCAGGCT C ATGGAATTGCTCTTTCTCCTCTGT | GGGG ACCACTTTGTACAAGAAAGCTGGGT C TGACCAAGGCCAGTTCGCAT |
| AT5G59960 | | protein of unkknown function | GGGG ACAAGTTTGTACAAAAAAGCAGGCT C ATGGAGAAAGTTTCGTCGCCG | GGGG ACCACTTTGTACAAGAAAGCTGGGT C GCTCTGCCTTGGCTTTTCTCG |
| AT1G11755 | LEW1 | LEAF WILTING 1 | GGGG ACAAGTTTGTACAAAAAAGCAGGCT C ATGGATTCGAATCAATCGATG | GGGG ACCACTTTGTACAAGAAAGCTGGGT C AGTTCCATAGTTTTGGTGGAC |
| AT5G48010 | THAS1 | THALIANOL SYNTHASE 1 | GGGG ACAAGTTTGTACAAAAAAGCAGGCT C ATGTGGAGGCTGAGAACTG | GGGG ACCACTTTGTACAAGAAAGCTGGGT C AGGGAGGAGACGTCGC |
| AT5G42600 | MRN1 | marneral synthase 1 | GGGG ACAAGTTTGTACAAAAAAGCAGGCT C ATGTGGAGACTGCGAATTGGAGC | GGGG ACCACTTTGTACAAGAAAGCTGGGT C AGAAACAAGCAGACGCAGAGC |
| AT4G38540 | MO2 | monooxygenase 2 | GGGG ACAAGTTTGTACAAAAAAGCAGGCT C ATGGAAGAAGAAGGCAGCCC | GGGG ACCACTTTGTACAAGAAAGCTGGGT C TGGGACAAGGCTTCCGC |
| AT1G17430 | | putative hydrolase | GGGG ACAAGTTTGTACAAAAAAGCAGGCT C ATGGCGTCCATGAAACACG | GGGG ACAAGTTTGTACAAAAAAGCAGGCT C AGCACTTAAGACAAACGAAG |
| AT5G01220 | SQD2 | sulfoquinovosyldiacylglycerol 2 | GGGG ACAAGTTTGTACAAAAAAGCAGGCT C ATGACGACTCTTTCTTCTATAAATC | GGGG ACCACTTTGTACAAGAAAGCTGGGT C CACGTTACCTTCCGGTACTGG |
| AT4G23430 | | putative short-chain dehydrogenase | GGGG ACAAGTTTGTACAAAAAAGCAGGCT C ATGTGGTTTTTTGGATCGAAAG | GGGG ACAAGTTTGTACAAAAAAGCAGGCT C AGAACTGCTTTCTCCTGATTG |
| AT5G15910 | | putative short-chain dehydrogenase | GGGG ACAAGTTTGTACAAAAAAGCAGGCT C ATGTTAAGGTCTCTGATTTGG | GGGG ACAAGTTTGTACAAAAAAGCAGGCT C GTGACCATGTTGAAGAATCC |
| AT1G72175 | | putative zinc finger protein | GGGG ACAAGTTTGTACAAAAAAGCAGGCT C ATGAATAGTCCACCGGAGAAC | GGGG ACCACTTTGTACAAGAAAGCTGGGT C CGAACCGGCATGACGGAAAT |
| AT1G25520 | PML4 | photosynthesis-affected mutant 71 like 4 | GGGG ACAAGTTTGTACAAAAAAGCAGGCT C ATGAGCTCGGTTTTGCAGGG | GGGG ACCACTTTGTACAAGAAAGCTGGGT C AGCCTCAACAGAAGTAAGATACG |
| AT3G23175 | | lesion inducing protein-related | GGGG ACAAGTTTGTACAAAAAAGCAGGCT C ATGAAGATGTCAATTAGCAAGGTC | GGGG ACCACTTTGTACAAGAAAGCTGGGT C TCCTTCGGTCTGCTTGTTCTTC |
| AT5G01750 | | unknown function | CTTTGTACAAAAAAGCAGGCT C ATGGAGCAGCCGTACGTGTAC | CTTTGTACAAGAAAGCTGGGT C GAGATAGTATACTTGGGCAACAATGGC |
| AT4G33360 | FLDH | farnesol dehydrogenase | GGGG ACAAGTTTGTACAAAAAAGCAGGCT C ATGGGCCCAAAGATGCCCAAC | GGGG ACCACTTTGTACAAGAAAGCTGGGT C GTAGTGAATGACGCCCAGACT |
| AT5G47990 | THAD | THALIAN-DIOL DESATURASE | GGGG ACAAGTTTGTACAAAAAAGCAGGCT C ATGGCATCAATGATCACTGTTGAC | GGGG ACCACTTTGTACAAGAAAGCTGGGT C AGTGTTTAGGTTTCGAGGAACAG |
| AT5G48000 | THAH | THALIANOL HYDROXYLASE | GGGG ACAAGTTTGTACAAAAAAGCAGGCT C ATGGATCGACGGGGCAAGTAT | GGGG ACCACTTTGTACAAGAAAGCTGGGT C GAGTGACTGGGAAATCTTGATAG |

**Table S3. List of microscopy settings for pollen tubes and root LDs.** All proteins were labeled with mCherry except for the ER marker ERD2 that was labeled with CFP. All LDs were stained with BODIPY 493/503.

| Microscopy of | 405 laser | 445 laser | 458 laser | 488 laser | 561 laser | 594 laser | detector "ERD-CF | detector BODIPY 49 | detector mCherry |
|---|---|---|---|---|---|---|---|---|---|
| At1g72175, At3g52730, At4g38540 | Y | N | N | Y | N | Y | 464-508 | 499-535 | 597-650 |
| At1g11755 | N | Y | N | Y | Y | N | 455-481 | 508-535 | 600-640 |
| At1g25520, At1g44170, At4g27760, At5g04070, At5g10050, At5g15860, At5g59960, At1g78800, At4g23430, At4g33180, At5g15910 | N | Y | N | N | Y | N | 455-481 | n.a. | 597-641 |
| At5g47990, At5g48000 | N | N | Y | N | Y | N | 463-552 | n.a. | 568-620 |
| At3g11430 | Y | N | N | Y | N | Y | 410-659 | 499-579 | 597-695 |
| At1g11755 | N | N | N | Y | Y | N | n.a. | 490-539 | 600-640 |
| At1g25520, At1g44170, At4g27760, At5g47990, At5g48000, At1g17430, At4g00550, At4g33180, At4g33360, At5g01220, At3g23175 | N | N | N | Y | Y | N | n.a. | 493-544 | 578-696 |
| At3g11430, At5g59960, At4g02370, At4g23430, At5g15910 | N | N | N | Y | Y | N | n.a. | 499-535 | 579-659 |
| At3g52730, At1g01610, At1g80950, At4g13160, At5g48010, At5g60620, At4g38540 | N | N | N | Y | Y | N | n.a. | 497-542 | 590-640 |
| At5g04070 | N | N | N | Y | Y | N | n.a. | 490-535 | 600-640 |
| At5g10050, At5g15860 | N | N | N | Y | Y | N | n.a. | 490-539 | 585-658 |
| At1g30130 | N | N | N | Y | Y | N | n.a. | 490-539 | 600-690 |
| At1g78800 | N | N | N | Y | Y | N | n.a. | 499-535 | 597-641 |
| roots | N | N | N | Y | N | N | n.a. | 495-550 | n.a. |
