## Supplementary material for "Arabidopsis root lipid droplets are hubs for membrane homeostasis under heat stress, and triterpenoid synthesis and storage": Figure S

### Supplemental Figures S1-S10

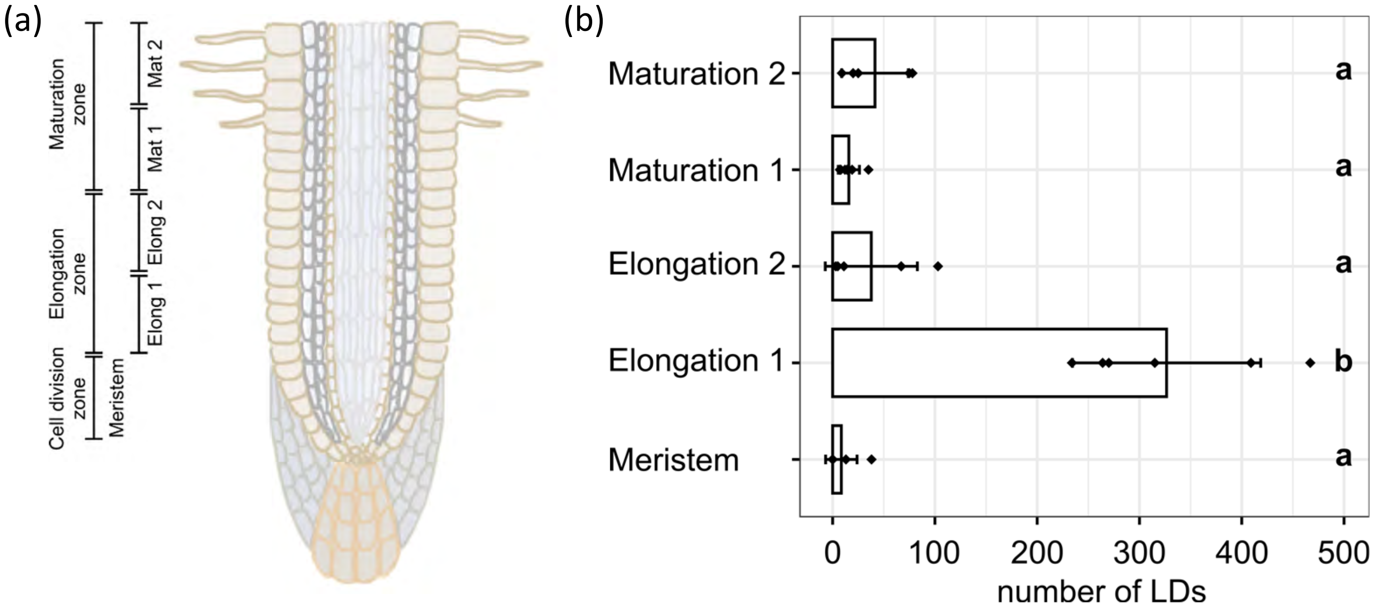


**Figure S1: LDs are enriched in different parts of the root elongation zone in Arabidopsis seedlings**. The roots of 7-day-old seedlings grown vertically on plates were fixated and stained with BODIPY 493/503. Plants were grown at 23°C prior to analysis. Median planes of different root zones (a) were imaged by CLSM (see Figure 1). For quantitative image analysis, areas up to 100 µm x 100 µm of each root micrograph were selected and the LDs within the selected areas were quantified in number and size using the particle analysis tool of ImageJ. Roots showed a significantly increased number of LDs in the elongation zone 1 compared to other regions of the root (b). Data was analyzed from n ≥ 5 individual roots. Plots display mean ± standard deviation. Data is identical to the data of the control presented in Figure 1.

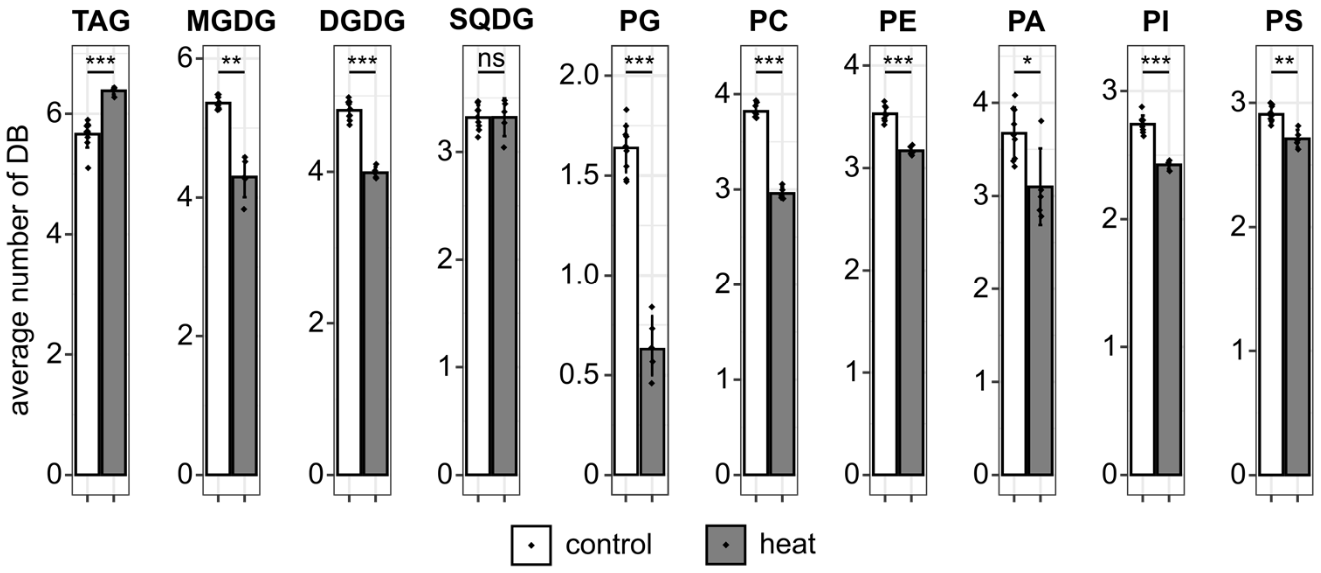


**Figure S2: The average number of double bonds decreases in most lipid classes in Arabidopsis seedlings subjected to heat stress**. Lipids were extracted from roots of 12-day-old seedlings grown vertically on plates. Plants were grown at 23°C, and in the case of heat stress moved for 24 h to 37°C prior to analysis. Lipids were analyzed by ESI-MS/MS. Determination of individual lipid species composition allowed the calculation of the average number of double bonds. Values are from n ≥ 5 biological replicates and are shown as mean ± standard deviation. Statistical differences were calculated by Welch's *t*-test using Benjamini-Hochberg correction for multiple comparisons and are represented as follows: $p > 0.05$ "ns", $p < 0.05$ "*", $p < 0.01$ "**", $p < 0.001$ "***".

DGDG, digalactosyldiacylglycerol; MGDG, monogalactosyldiacylglycerol; PA, phosphatidic acid; PC, phosphatidylcholine, PE, phosphatidylethanolamine; PG, phosphatidylglycerol; PI, phosphatidylinositol; PS, phosphatidylserine; sulfoquinovosyldiacylglycerol; TAG, triacylglycerol.

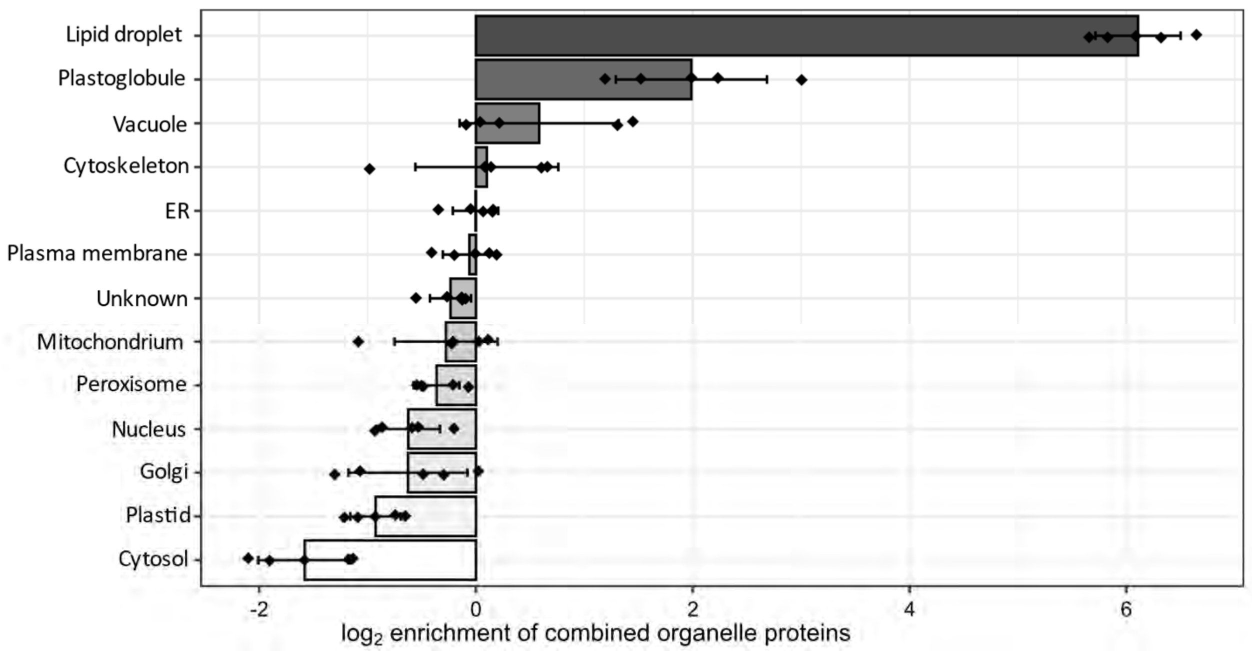


**Figure S3: Based on the overall proteomic data, no other organelles are strongly enriched in the LD-enriched fraction.** Roots of *tgd1-1 sdp1-4* mutant plants were grown in axenic root culture and an LD-enriched fraction isolated. The proteome of this fraction and a total protein fraction was investigated by LC-MS/MS and relative iBAQ values were calculated. To analyze enrichments of different subcellular organelles, proteins were assigned to their subcellular localization according to the online Plant Proteome Database and previous reports of known plant LD proteins. Only proteins identified by at least two peptides and present in all replicates of at least one fraction were used for this analysis. Furthermore, assignment to organelles was only done for proteins with a unique localization, all other proteins were designated as “unknown”. The combined protein abundance of all marker proteins of the different organelles in the LD-fraction was then normalized to the respective combined protein abundances in total extract samples and the resulting enrichment ratio was $log_2$-transformed. n = 5 biological replicates.

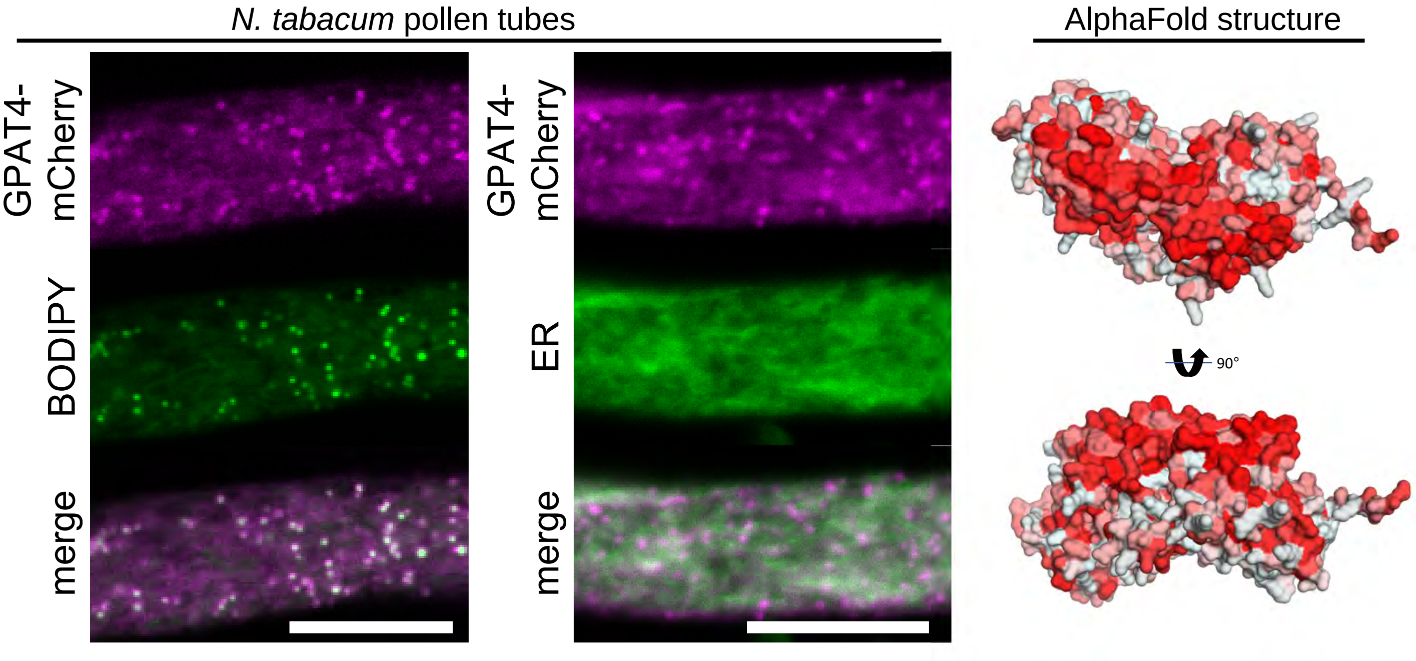


**Figure S4: Subcellular localization of Arabidopsis GLYCEROL-3-PHOSPHATE ACYLTRANSFERASE 4 (GPAT4) in *N. tabacum* pollen tubes.** mCherry-tagged GPAT4 was expressed in *N. tabacum* pollen tubes. Either LDs were stained with BODIPY 493/503 or the ER marker ERD2-CFP was co-expressed. Images are single planes obtained by CLSM. GPAT4 clearly colocalized with LDs in pollen tubes. Each image is representative for 10 pollen tubes. Bars, 10 µm. As shown on the right, the protein structure of GPAT4, as predicted by AlphaFold2, shows a hydrophobic surface on one side of the protein.

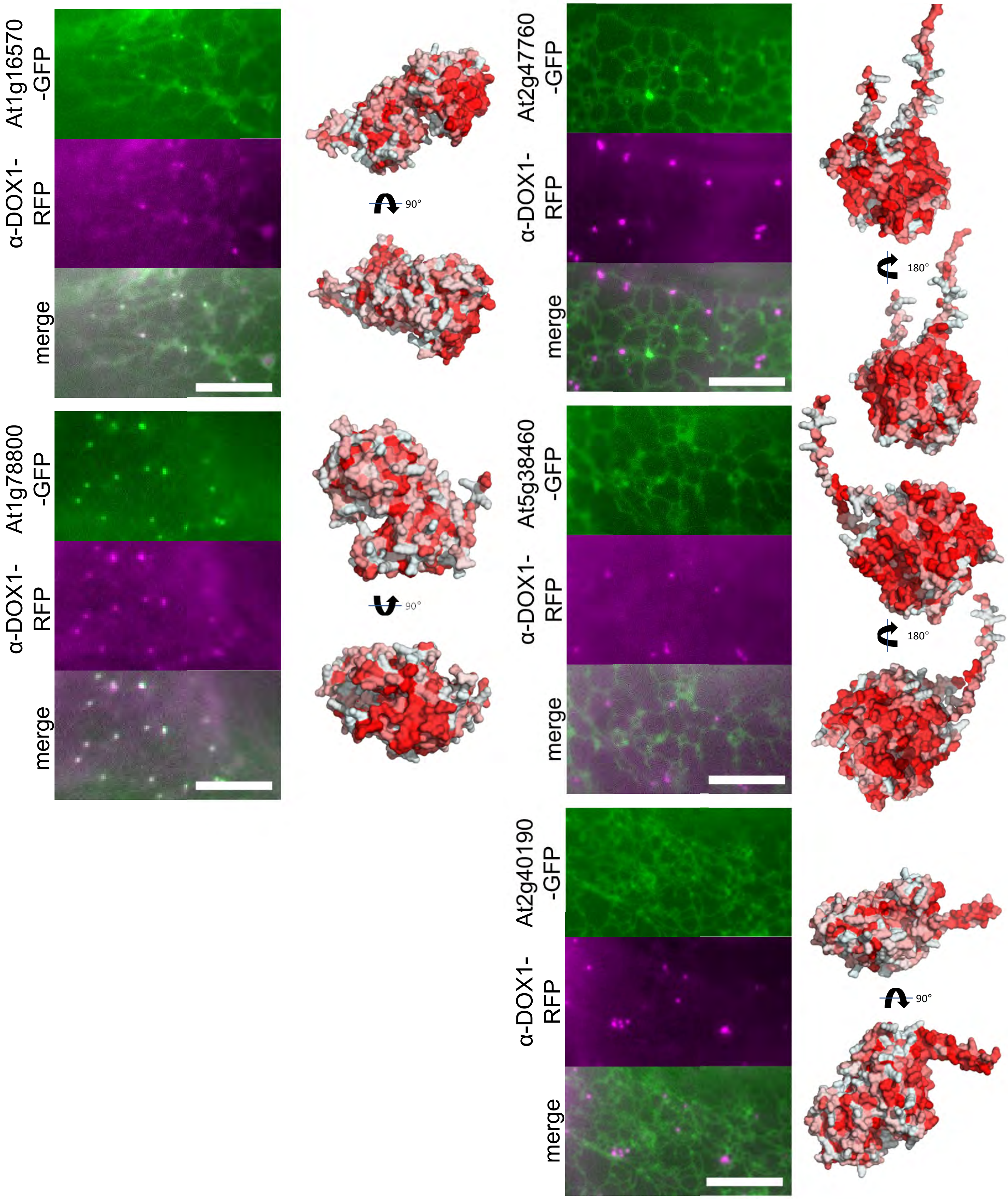


**Figure S5: Subcellular localization of Arabidopsis N-glycan biosynthetic enzymes in *N. benthamiana* leaves.** Indicated GFP-tagged proteins were expressed in *N. benthamiana* leaves. The formation of LDs was induced by heat stress and α-DOX1-RFP was co-expressed, serving as an LD marker protein. Images were obtained by fluorescence microscope. At1g16570 and At1g78800 colocalize with the LD marker, while the other three members of the protein family display a reticular pattern. Each image is representative for at least 4 leaf areas. Bars, 10 µm. As shown on the right, the protein structures of At1g16570 and At1g78800, as predicted by AlphaFold2, show a hydrophobic surface on one side of the protein.

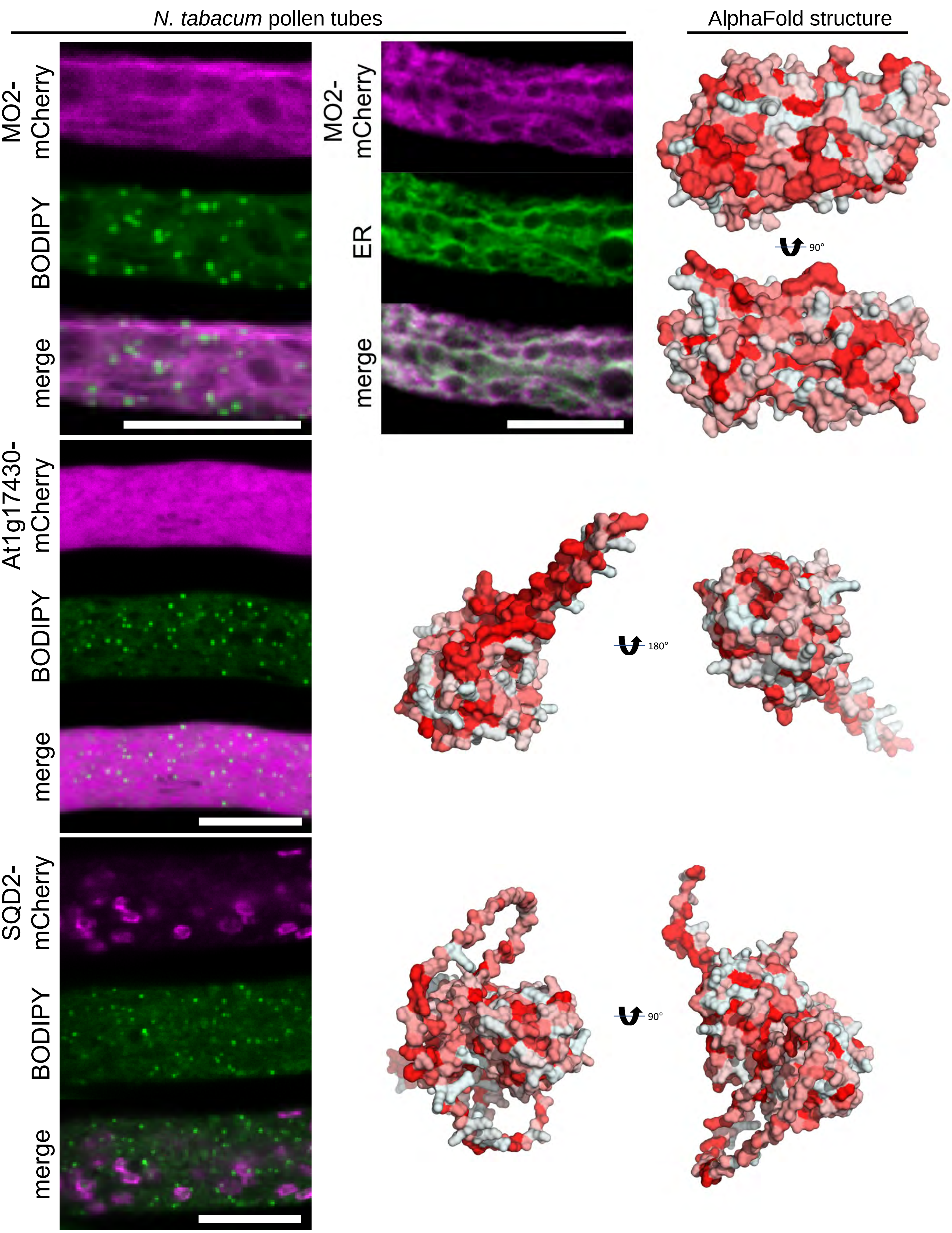


**Figure S6: Subcellular localization of selected Arabidopsis candidate root LD proteins in *N. tabacum* pollen tubes.** Indicated mCherry-tagged proteins were expressed in *N. tabacum* pollen tubes. Either LDs were stained with BODIPY 493/503 or ERD2-CFP was co-expressed, serving as an ER marker. Images are single planes obtained by CLSM. MONOOXYGENASE 2 (MO2) colocalized with the ER marker while the putative hydrolase At1g17430 displayed cytosolic localization. SULFOQUINOVOSYLDIACYLGLYCEROL 2 (SQD2) localizes to structures with high resemblance to plastids in pollen tubes. Each image is representative for at least 7 pollen tubes. Bars, 10 µm. The top two structures, as predicted by AlphaFold2, show hydrophobic surface regions but not a flat face like involved in several LD-binding proteins. SQD2 does not display any larger hydrophobic regions.

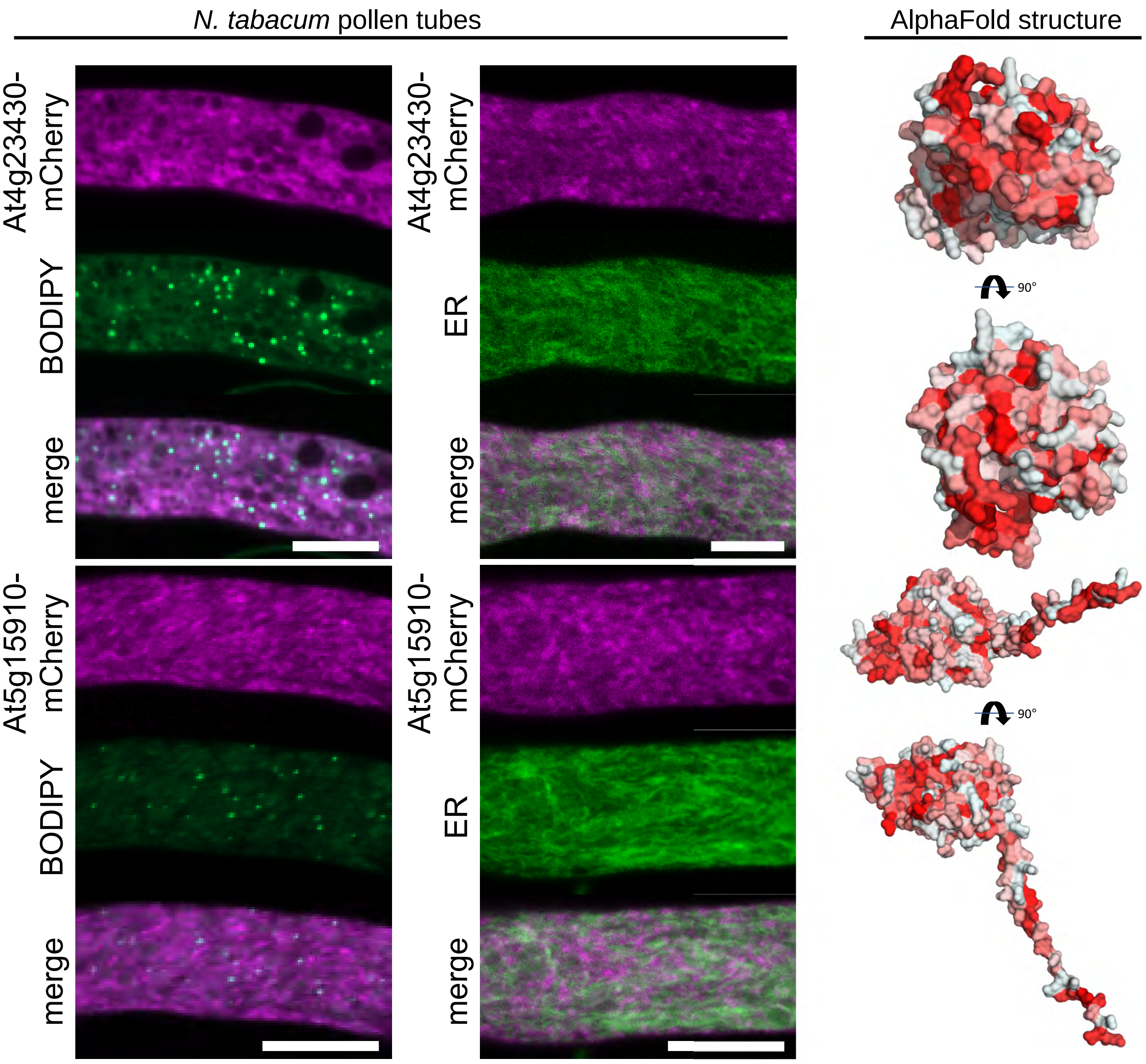


**Figure S7: Subcellular localization of Arabidopsis putative dehydrogenases in *N. tabacum* pollen tubes.** Indicated mCherry-tagged proteins were expressed in *N. tabacum* pollen tubes. Either LDs were stained with BODIPY 493/503 or ERD2-CFP was co-expressed, serving as an ER marker protein. Images are single planes obtained by CLSM. Both putative dehydrogenases did not colocalize with LDs or the ER. Each image is representative for 6 pollen tubes. Bars, 10 µm. As shown on the right, the structures of both proteins, as predicted by AlphaFold2, show hydrophobic surface regions, but not a flat face as found in several LD-binding proteins.

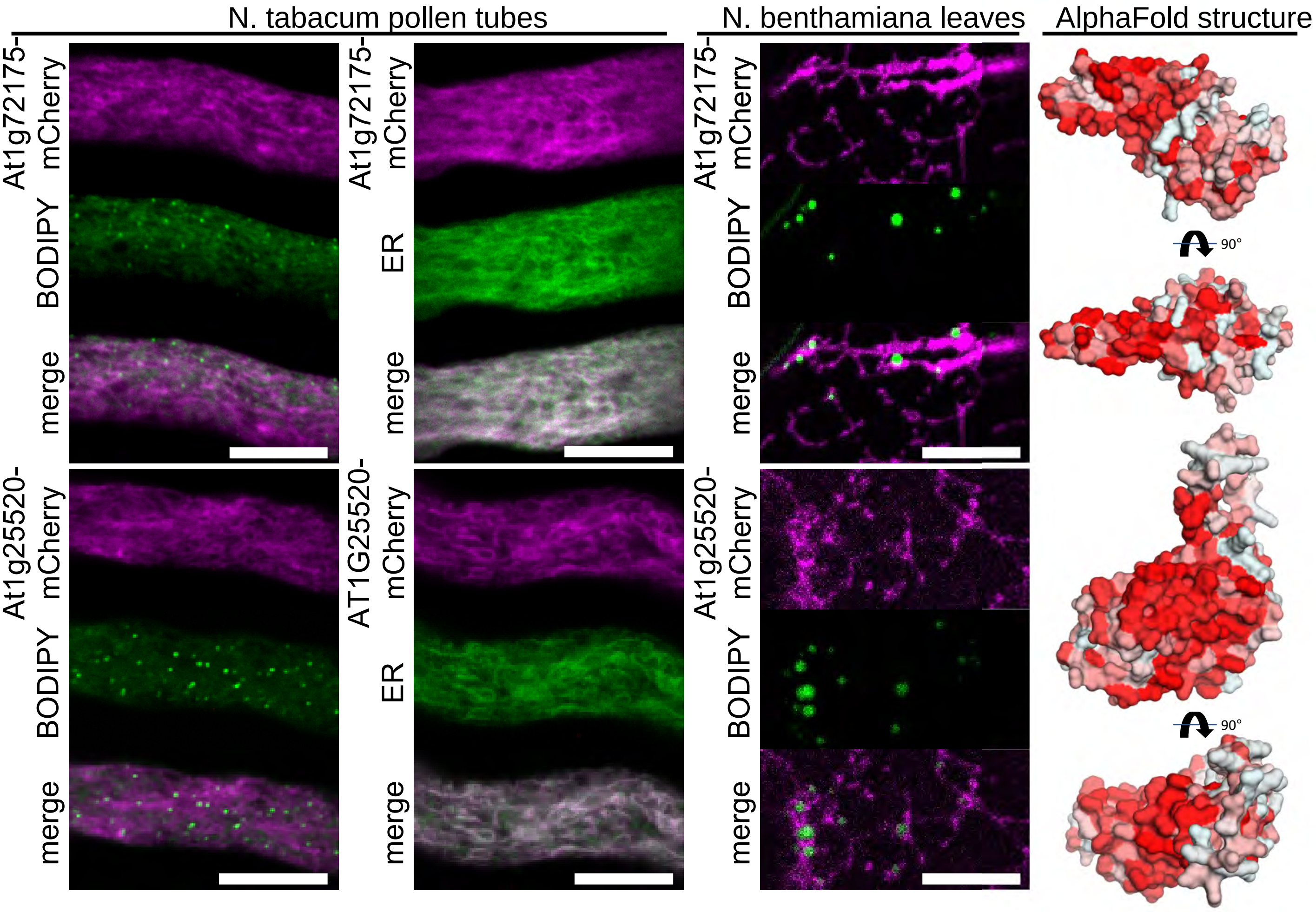


**Figure S8: Subcellular localization of selected candidate Arabidopsis root LDs proteins with unknown function in *N. tabacum* pollen tubes and *N. benthamiana* leaf cells.** Indicated mCherry-tagged proteins were expressed in either *N. tabacum* pollen tubes or *N. benthamiana* leaves. LDs were stained with BODIPY 493/503 or ERD2-CFP was co-expressed, serving as an ER marker protein. Images are single planes obtained by CLSM. Both the putative zinc finger protein (At1g72175) and a protein of unknown function (PHOTOSYNTHESIS-AFFECTED MUTANT 71 LIKE 4, At1g25520) targeted the ER in pollen tubes and reticular structures in leaves. Each image is representative for at least 9 pollen tubes or 4 leaf areas. Bars, 10 µm. As shown on the right, the structures of both proteins, as predicted by AlphaFold2, show hydrophobic surface regions, but not a flat face as found in several LD-binding proteins.

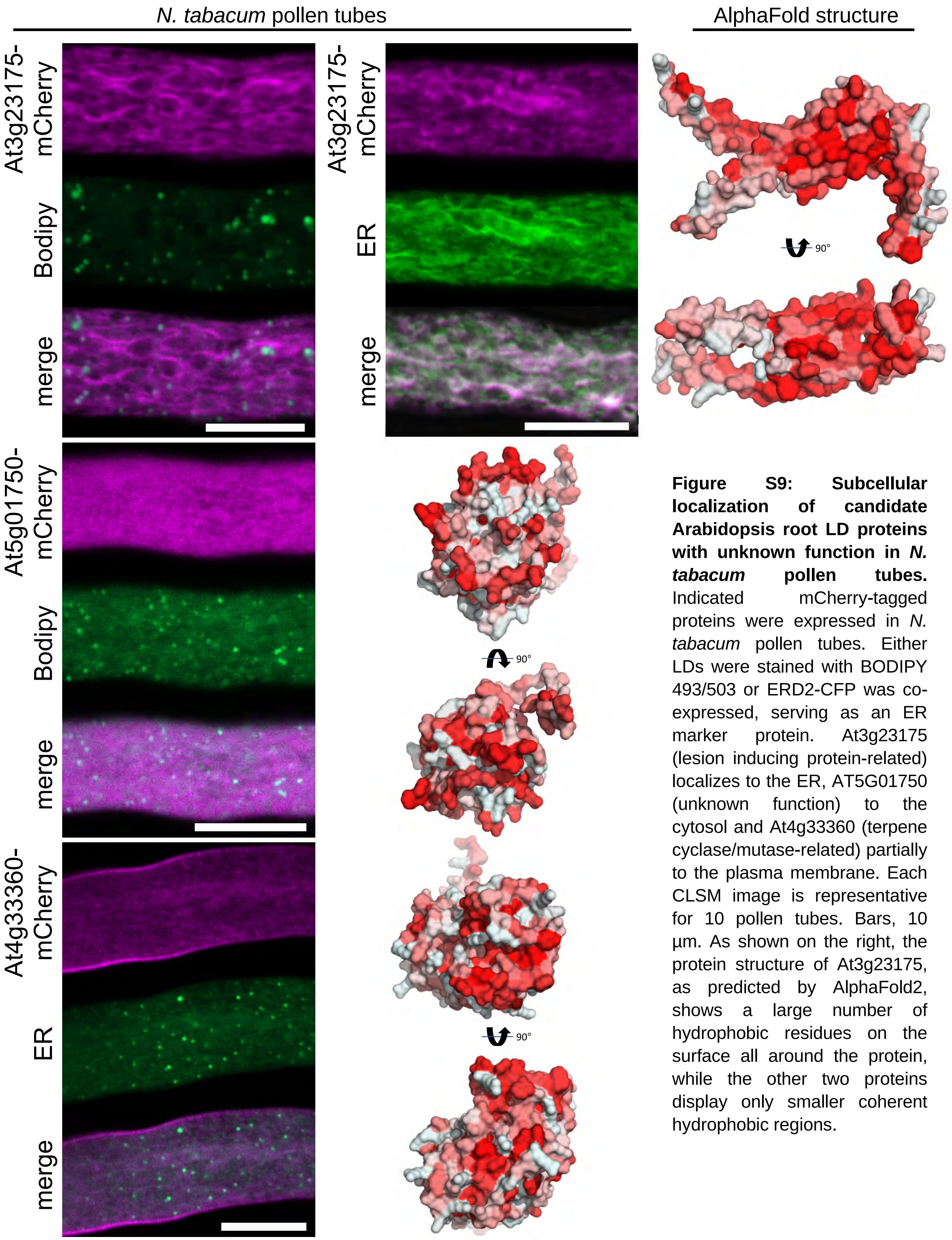


**Figure S9: Subcellular localization of candidate Arabidopsis root LD proteins with unknown function in *N. tabacum* pollen tubes.** Indicated mCherry-tagged proteins were expressed in *N. tabacum* pollen tubes. Either LDs were stained with BODIPY 493/503 or ERD2-CFP was co-expressed, serving as an ER marker protein. At3g23175 (lesion inducing protein-related) localizes to the ER, AT5G01750 (unknown function) to the cytosol and At4g33360 (terpene cyclase/mutase-related) partially to the plasma membrane. Each CLSM image is representative for 10 pollen tubes. Bars, 10 µm. As shown on the right, the protein structure of At3g23175, as predicted by AlphaFold2, shows a large number of hydrophobic residues on the surface all around the protein, while the other two proteins display only smaller coherent hydrophobic regions.

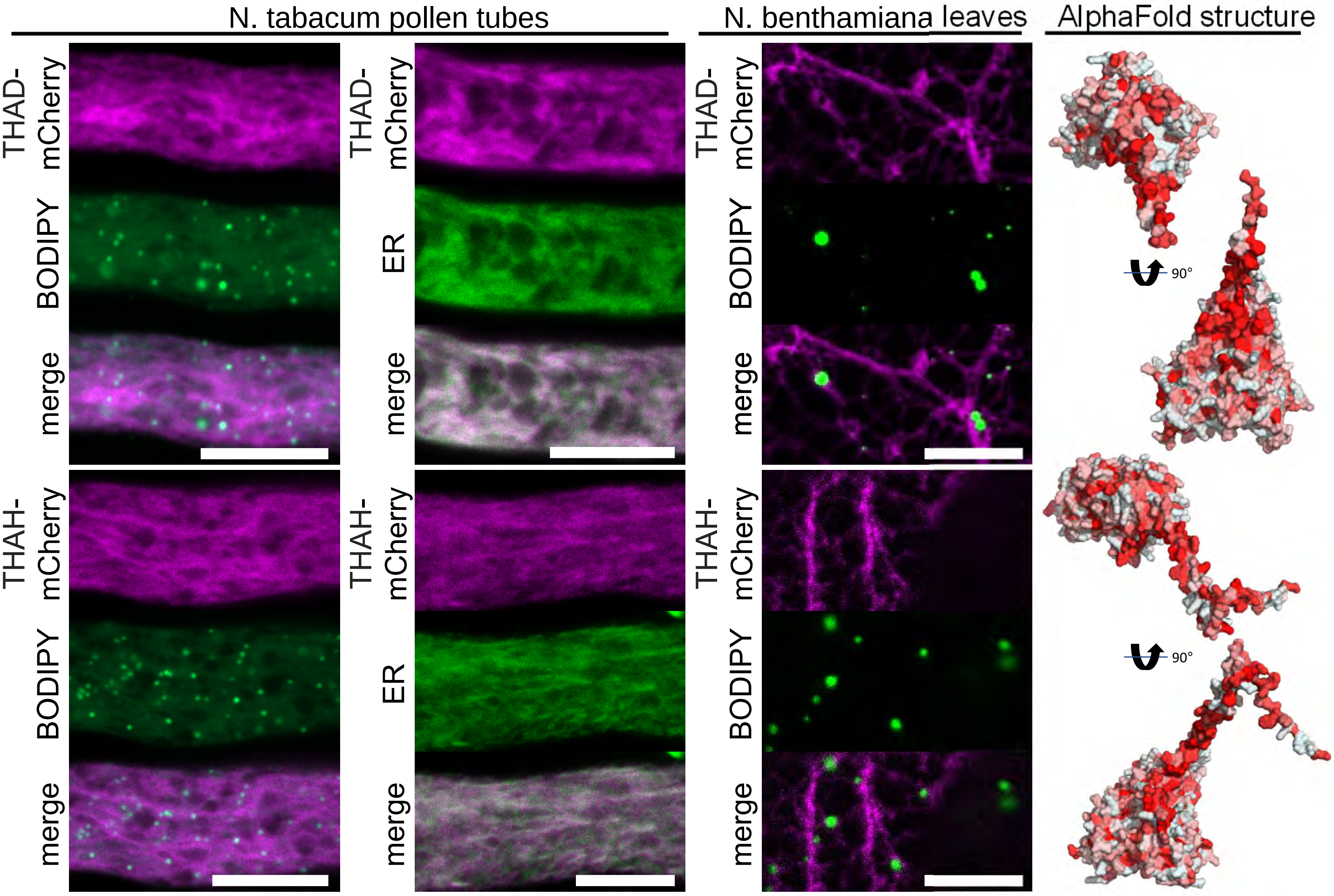


**Figure S10: Subcellular localization of various Arabidopsis enzymes acting downstream of thalianol synthase.** Indicated mCherry-tagged proteins were expressed in *N. tabacum* pollen tubes or *N. benthamiana* leaves. Either LDs were stained with BODIPY 493/503 or ERD2-CFP was co-expressed, serving as an ER marker protein. Images are single planes obtained by confocal microscopy. Both THALIAN-DIOL DESATURASE (THAD) and THALIANOL HYDROXYLASE (THAD) targeted the ER in pollen tubes and reticular structures in leaves. Each CSLM image is representative for at least 10 pollen tubes or 4 leaf areas. Bars, 10 µm. As shown on the right, the structures of both proteins , as predicted by AlphaFold2, show helical or rod-like hydrophobic regions but no flat hydrophobic surface areas.
